# Pre-Sensory Spontaneous Activity Accelerates Coordinated Maturation of Synaptic Partners and Drives Transition to the Mature Physiological Phenotype

**DOI:** 10.1101/2025.08.28.672917

**Authors:** Daniel T. Heller, Nikollas M. Benites, Emily M. Amick, Andre Dagostin, Samuel M. Young, Henrique von Gersdorff, George A. Spirou

**Author notes:** Corresponding author: George A. Spirou University of South Florida, 4202 E. Fowler Avenue, ENG 030, Tampa, FL 33620-9951. Equal contribution (Daniel T. Heller, Nikollas M. Benites). **Competing Interest Statement:** George Spirou, Ph.D., is a co-founder and has a financial interest in the software syGlass.

## Abstract

Pre-sensory (ps), peripherally-generated spontaneous activity (SA) is important for establishing basic topography of central components of sensory pathways. However, roles for psSA in sculpting connections at the single neuron level and in driving functional maturation of postsynaptic targets are difficult to isolate and therefore little explored. We capitalized on the temporal onset of cochlea-generated psSA just prior to growth of the calyx of Held (CH), the largest nerve terminal in the mammalian brain, and its well-defined circuit topography to explore the causal roles for psSA in targeted synaptogenesis and postsynaptic functional maturation. To this end, we developed a viral vector to rapidly express tetanus neurotoxin (TeNT), which blocks neurotransmission, and selectively introduced it into CH-forming neurons at the onset of psSA. TeNT expression did not prevent but delayed and altered the developmental trajectory of neural circuit maturation, which in sum prevented the postsynaptic neuron from acquiring its mature phasic firing phenotype. Thus, psSA initiates and drives circuit maturation along central sensory pathways, but alternative mechanisms can partially compensate in its absence.

## Introduction

The fundamental aspects of mature neuronal circuit organization are orchestrated through a combination of epigenetic and transcriptional regulation, molecular guidance cues and spontaneous activity (SA)^1^. In mammals, SA in several sensory organs initiates at pre-sensory (ps) ages and has a characteristic bursting pattern^2–5^. Silencing of psSA in primary afferent neurons disrupts topographic sorting of axons in the CNS, such as formation of ocular dominance columns, glomerular targeting by olfactory sensory neurons, and organization of mystacial whisker barrelettes and barreloids^6–9^. However, manipulations that silence neurotransmission genetically overlap embryonic events of cell migration and axon targeting, preceding bursting psSA^9,10^ and later manipulations have focused on formation of gross topography for innervation^2–5^. Hence, little is known at the single neuron level and with high temporal precision about how psSA, from its onset, regulates circuit refinement and coordinated maturation of pre- and postsynaptic partners.

In the auditory system, the onset of cochlea-driven psSA occurs just after birth^11,12^ when cell migration and initial axon targeting are complete^13^, and just prior to the rapid growth of the calyx of Held (CH), which forms synaptic contact between second and third order neurons in the brainstem^14–17^. Therefore, we capitalized on this temporal arrangement and rapid establishment of circuit topography during a pre-sensory period^15–17^ that ends with opening of the ear canals beginning at P10^18^, to hypothesize that psSA drives coordinated pre- and postsynaptic maturation of the CH and its synaptic partner, called a principal neuron (PN). However, genetic manipulations which eliminated or downregulated psSA in the cochlea have resulted in minimal deficits to CH formation or synaptic transmission^19–22^. Notably, peripheral afferent neurons can compensate homeostatically and become hyperexcitable to preserve SA^23^. Neurons along central pathways leading to primary sensory cortex in other systems may compensate similarly.

Among strategies to study effects of psSA at central locations, expression of tetanus neurotoxin (TeNT) prevents AP-induced neurotransmission while preserving AP generation in the afferent axon, thereby isolating specific roles for synaptic communication^9,24–27^. To this end, we created a Helper-dependent Adenoviral vector which contained a transgene expression cassette^28^ for rapid onset of TeNT expression^29^ to directly transduce CH-forming neurons in the cochlear nucleus. Virus injection on the day of birth (postnatal day (P)0) led to expression by P2, as bursting activity is initiating^11,12^ and prior to growth of the CH and topographic refinement of innervation^15,16,30^. In contrast to a lack of effect on CH structure with TeNT expression after CH formation^26^, early TeNT expression delayed CH formation, and CHs and PNs were smaller in volume. Additionally, we found maturation of biophysical and AP properties of PNs was delayed and/or attenuated, which in sum prevented the characteristic transition from tonic to mature phasic firing phenotype. Computational modeling of electrophysiological data indicated a psSA-driven program to balance voltage-gated K^+^ channel expression during functional maturation of PNs. Thus, targeted silencing of synaptic transmission at a single synaptic station revealed roles for psSA to activate maturational programs, that neurotransmission-independent mechanisms could also initiate after a delay, and the necessary role for psSA to drive transition to the mature physiological phenotype of the postsynaptic neuron.

## Results

### Viral Vector Expression of TeNT is Rapid Following Injection at P0 and Delays Growth of the CH

To directly test the role of pre-sensory (ps) spontaneous activity (SA) in maturation of the calyx of Held (CH) and its postsynaptic neuron, the principal neuron (PN) of the medial nucleus of the trapezoid body (MNTB), we developed a Helper-dependent Adenoviral (HdAd) vector to transduce the presynaptic neuron that gives rise to the CH, called a globular bushy cell (GBC), with the light chain subunit of tetanus neurotoxin (TeNT) (Fig. 1A and B). The light chain of TeNT mediates the proteolytic cleavage of the synaptic vesicle fusion protein synaptobrevin-2, also known as vesicle-associated membrane protein 2 (VAMP2) (Fig. 1A)^29^, preventing opening of a fusion pore and subsequent neurotransmitter release^31^. The HdAd construct independently co-expresses the TeNT light chain driven by the overexpression pUNISHER cassette allowing for rapid expression^28^, along with a fluorescent reporter molecule (mCherry) independently driven by the 470-bp human synapsin promoter. Unilateral virus injections allowed for an in-slice control comparing transduced and non-transduced calyces in the MNTB contralateral and ipsilateral to the injection site, respectively (Fig. 1C-C2). Previously we have shown pUNISHER driven expression of EGFP exhibited normal synaptic transmission and structural development of the CH^28^. mCherry expression in GBCs was detectable within 48 hours (Fig. S1A) and labeled their cell bodies and axonal projections along the ventral brainstem surface, which crossed the midline to form synapses onto PNs of the contralateral (c)MNTB (Figs. 1B, 1C-C2, and S1). Thus, TeNT expression at the CH:PN synaptic connection occurred coincident with the onset of cochlea induced bursting activity and prior to initial growth and topographic refinement of the CH^11,12,16,17,32^. We found that a small number of cell bodies in the contralateral ventral cochlear nucleus were trans-synaptically labeled (Figs. 1C, S1), whose commissural projections primarily innervate multipolar cells and spherical bushy cells^33,34^. The few transduced calyces in the iMNTB (Figs. 1C1, S1C and Table S1) likely reflect the ≈5% ipsilateral projecting GBCs^35^. We did not observe retrogradely labeled fibers in the auditory nerve, so we conclude that SA generation in the spiral ganglion remained intact.

**Figure 1.**
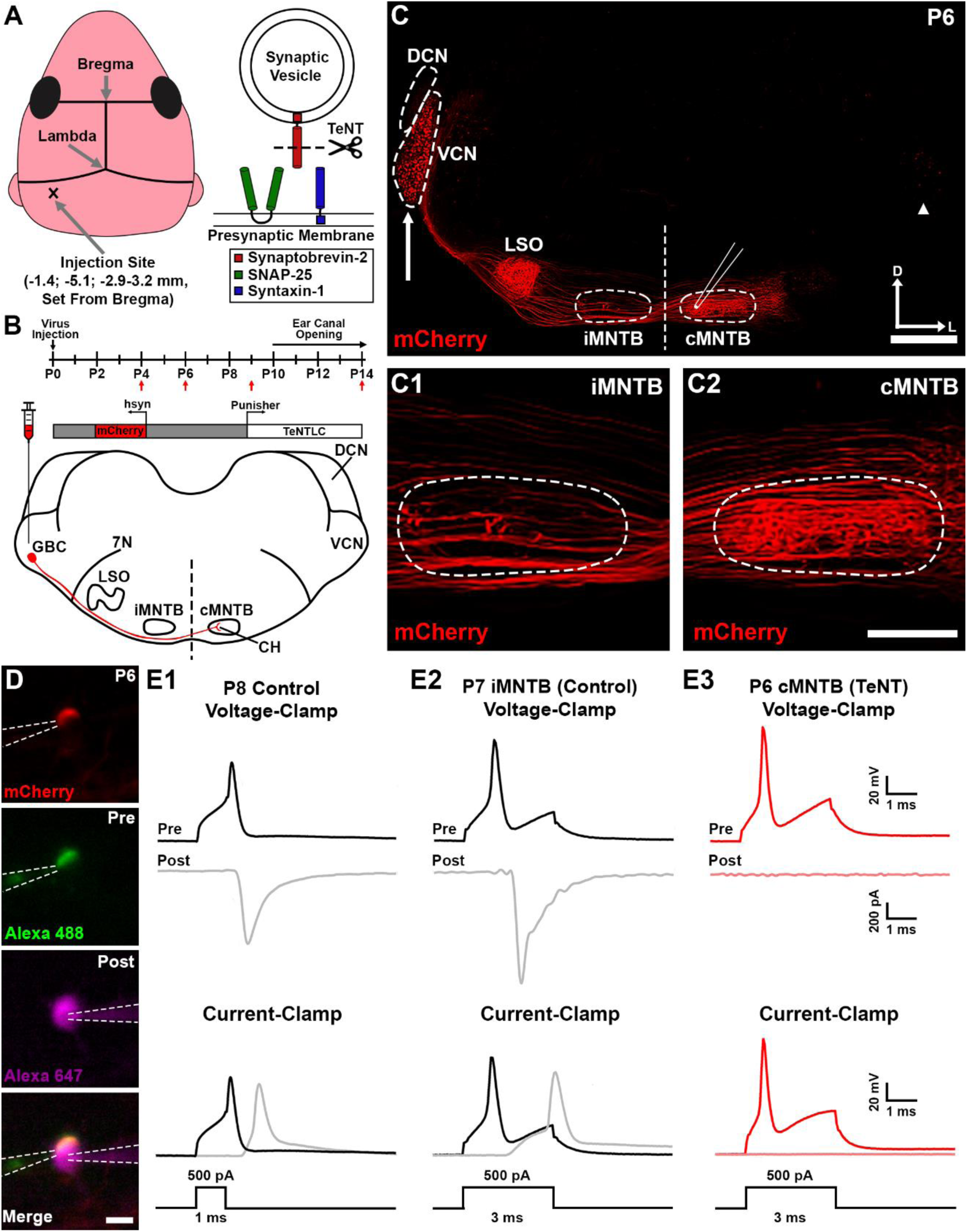
Experimental protocol and efficacy, shown at P6-P8, of unilateral viral vector injections at P0 into the VCN expressing TeNT to block synaptic activity. (A), Schematic showing the injection coordinates for a P0 mouse pup and the mechanism of action for TeNT light chain cleaving the synaptic vesicle fusion protein Synaptobrevin-2. (B), (Top) Timeline for viral vector injection and experimental data collection (red arrows). (Bottom) Schematic of auditory brainstem circuit, whereby virus injection into ventral cochlear nucleus (VCN) transduces globular bushy cells (GBCs), which form large calyx of Held (CH) terminals onto PNs in contralateral medial nucleus of the trapezoid body (cMNTB). HdAd construct expresses tetanus neurotoxin (TeNT) light chain driven by the Punisher overexpression cassette and mCherry driven by 470 bp human synapsin (hsyn) promoter. (C), High titer viral vector injection into VCN (arrow) at P0 labels GBCs whose axons travel along ventral brain surface, cross midline (dashed line) and terminate in cMNTB; small number of GBCs innervate iMNTB. Pipette outline indicates recordings from medial 1/3 of MNTB. Other VCN neurons are transduced and innervate lateral superior olive (LSO) and dorsal cochlear nucleus (DCN). Some cells are anterogradely labeled in non-injected VCN. Dorsal and lateral axes are indicated above scale bar. Scale bar: 500 μm. (C1 and C2), High-magnification images of iMNTB and cMNTB. Scale bar: 200 μm. (D), Paired recordings of transduced CH (mCherry, double-labeled with Alexa 488 conjugated dextran in pipette; Merge panel at bottom) and postsynaptic principal neuron (PN; labeled with Alexa 647 conjugated dextran). Dashed lines outline pre- and postsynaptic patch pipettes. Scale bar: 10 μm. (E1-E3), Paired recordings in current-clamp (CH) and voltage and current-clamp (PN) for non-injected (control: E1) and from iMNTB (non-transduced control; E2) and cMNTB (TeNT expression; E3) following viral injection. Evoked responses in PNs, following a 1-3 ms, 500 pA depolarizing current injection via the presynaptic recording pipette, in voltage (EPSC, top) and current-clamp (AP, bottom) configuration show TeNT abolished neurotransmission. The experimental recordings in the cMNTB (E3) correspond to the fluorescence images in panel (D).

### AP-Evoked Synaptic Transmission is Abolished at the Calyx of Held Following *In Vivo* Expression of TeNT

Since GBCs innervate PNs from embryonic day (E)17 via small terminals^17,30,36^, we hypothesized that rapid expression of TeNT would permit continued innervation of the PN but prevent growth of the CH. Interestingly, immunolabeling for vesicular glutamate transporters 1 and 2 (Vglut1/2) revealed that CHs did grow onto PNs (Table S1). However, at P4 only a small fraction of PNs in the cMNTB were contacted by CHs (∼22%) relative to the ipsilateral (i)MNTB (∼90%), the latter of which was consistent with previous ultrastructural analysis^16^. By P6 most PNs in the cMNTB were innervated by CH terminals (∼82%; Table S1). Thus, TeNT exerted rapid effects and delayed CH growth by about two days.

Since CH growth persisted following TeNT expression, we took advantage of the giant size of the terminal to evaluate synaptic transmission via simultaneous paired pre- and postsynaptic whole-cell recordings in *ex vivo* brain slices at P6-P8 (Fig. 1D and E1-E3)^37,38^. Alexa fluorophores of two different wavelengths (488 and 647; 50 μM) were incorporated in the patch pipettes for visual confirmation of simultaneous presynaptic (mCherry from HdAD and Alexa 488 co-labeling) and postsynaptic (Alexa 647) recordings (Fig. 1D). In all paired recordings from the cMNTB (n = 4) a depolarizing current injection in transduced calyces (TeNT expression) elicited a presynaptic action potential (AP), indicating that TeNT did not alter psSA upstream of the CH:PN synapse (Figs. 1E3, S2B-D). However, in all cell pairs the presynaptic AP failed to evoke an EPSC or AP in the postsynaptic PN, measured in voltage and current clamp, respectively (Figs. 1E3, S2B-D). To ensure paired recordings were performed on healthy PNs, spontaneous EPSCs (sEPSCs) and the firing pattern from postsynaptic depolarizing current injections were monitored with consistent results to other experimental recordings (Fig. S2A1-A2, B1-D1, and B2-D2). For comparison, recordings were performed on P6-P8 CH:PN synaptic pairs in the iMNTB, following viral injection, or in non-injected mice. All paired recordings from these control conditions (iMNTB, n = 3; non-injected, n = 2) displayed the characteristic evoked response (EPSC in voltage-clamp and AP in current-clamp) in the PN following a presynaptic spike in the CH (Fig. 1E1 and E2).

### Spontaneous Neurotransmission (sEPSC) is Greatly Reduced at the Calyx of Held Following *In Vivo* Expression of TeNT

Prior to the growth phase of the CH (after P2), MNTB PNs are hyperexcitable, whereby spontaneous vesicle release is likely sufficient to trigger an AP^30,39^. Previous reports utilizing TeNT to block synaptic neurotransmission showed a reduction of ∼70-90% in sEPSCs^24,26,27^. To more completely assess effects of TeNT, we recorded sEPSCs from PNs in both cMNTB and iMNTB following unilateral virus injection, and from non-injected control animals in order to assay its effect on spontaneous neurotransmission prior to (P4, P6, and P9) ear canal opening at P10, and P14, soon after mice are sensitive to low threshold sound^18^. Although direct tissue stimulation can activate the cochlea at high intensity from about P7 and air-conducted sound at higher intensity^40,41^, neural responses *in vivo* are likely dominated by cochlea-generated SA until P10^11,12,42^. In parallel with eyelid opening at about the same age^43^, we refer to ages until P9 as pre-sensory. Representative recordings at P6 reveals a significant reduction in the frequency and a slower time course following TeNT expression compared to PNs recorded from the iMNTB and non-injected mice (Fig. 2A). This reduced frequency was consistent from P4 to P9 (Fig. 2B and Table S2), whereby ∼30% of neurons had a rate ≤ 0.3 Hz, and increased slightly (all cells ≤ 2.5 Hz) after ear canal opening (P14). The residual low frequency of TeNT resistant events could be mediated by non-canonical SNARE proteins, VAMP7 (also known as TeNT-insensitive VAMP) and Vps10p-tail-interactor-1a (Vti1a)^44,45^ at the CH or reflect non-CH synaptic inputs^46^. iMNTB and non-injected controls had similarly greater sEPSC frequencies, except at P9 and P14, when iMNTB PNs showed a reduced frequency more similar to cMNTB (Fig. 2B). This decreased frequency in the iMNTB may represent homeostatic compensation emergent in the neural circuit. At P14, ear canal opening may contribute sensory-driven compensation in the neural circuit.

**Figure 2.**
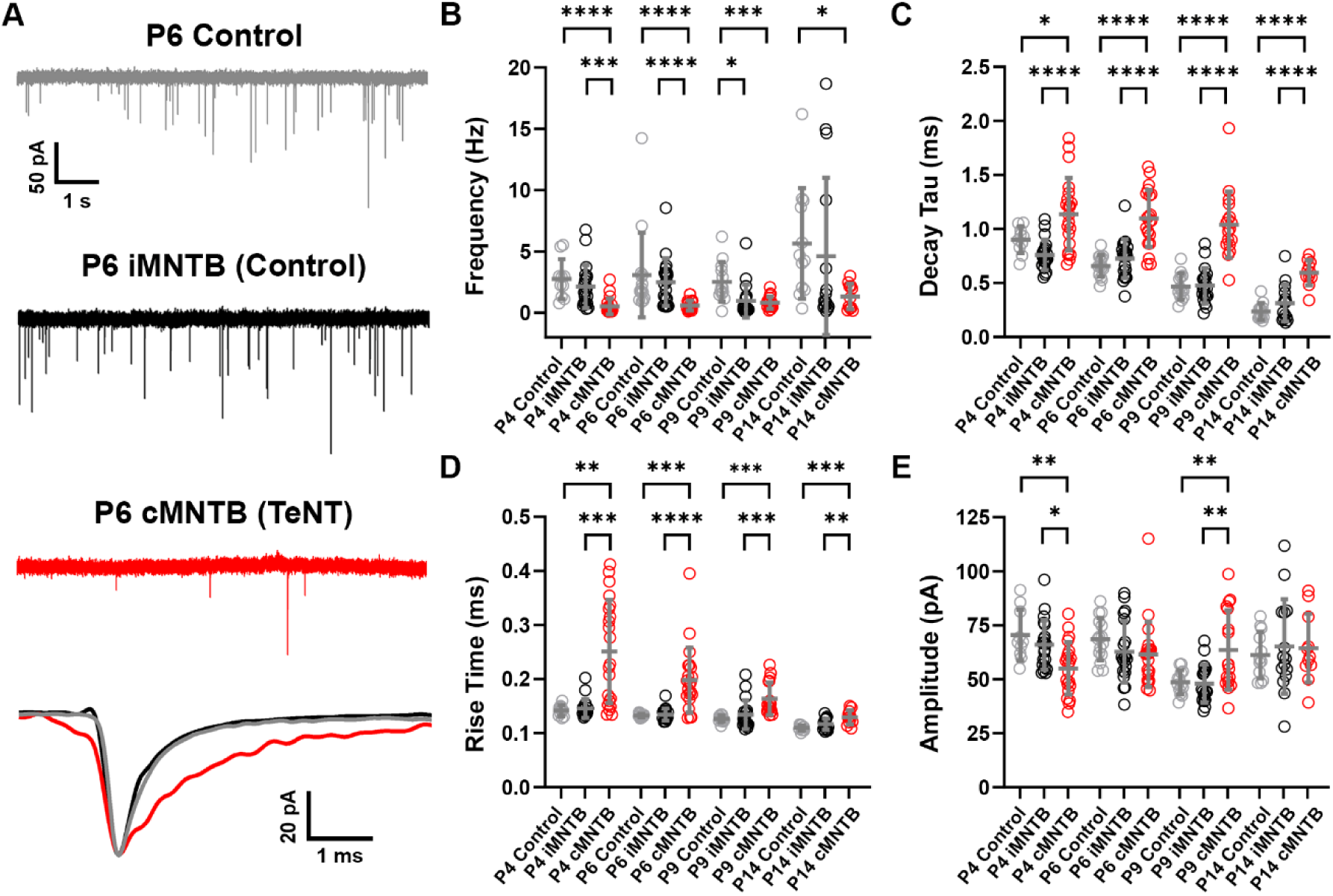
TeNT reduces frequency and slows kinetics of spontaneous (s)EPSC in MNTB PNs. (A), Exemplary voltage-clamp traces of P6 PN sEPSCs from non-injected control mouse (grey traces) and from the iMNTB (non-transduced control, black traces) and cMNTB (transduced, red traces) of unilaterally injected animals. Cells clamped at holding potential of −73 mV. Recordings were made after bath application of Gabazine (GABA_A_ receptor antagonist, 10 μM) and strychnine (glycine receptor antagonist, 2 μM). Averaged sEPSC waveforms from each cell shown at bottom with expanded time scale and color-coordinated. (B-E), Plots show reduced frequency, increased average decay time constant (τ) and rise time, and near constant sEPSC amplitude in cMNTB PNs across age. Non-injected control recordings from MNTB PNs (grey circles; P4, n = 10; P6, n =15; P9, n = 13; P14, n = 12) were compared to PNs in the iMNTB (black circles, Control; P4, n = 20; P6, n =20; P9, n = 19; P14, n = 15) and cMNTB (red circles, TeNT; P4, n = 21; P6, n =21; P9, n = 17; P14, n = 11) following unilateral viral injection. Mean (horizontal gray lines) and standard deviation (error bars). Significance levels are indicated according to the convention: *P < 0.05, **P < 0.01, ***P < 0.001, ****P < 0.0001, and not significant (ns). Statistical results for panels (B-E) are reported in Table S6.

sEPSCs in cMNTB PNs showed consistently slower decay times from P4-P14, and a delayed (5-7 days) and slower rate of decrease in these values from P4-P9 relative to iMNTB and non-injected controls (Fig. 2C and Table S2). Both non-injected mice and the iMNTB of injected mice had similar values across postnatal development. sEPSC rise times were also generally slower in the experimental group with a delayed decrease (5 days) from P4-P14, but due to the small average values and the variance within groups, statistical significance was less consistent (Fig. 2D). sEPSC amplitudes in cMNTB were unchanged from P4-P14, but both control groups declined from P4-P9, then increased at P14 to match cMNTB values (Fig. 2E). Both control groups showed no difference across metrics and age, with the exception of the frequency at P9 (Fig. 2B), confirming the utility of our unilateral viral injection approach with an in-slice iMNTB control (Fig. 1A-C). Overall, viral vector-mediated TeNT expression eliminated AP-evoked and greatly reduced spontaneous vesicle release during the critical growth period of the CH^16,30,32,39^.

### Pre-Sensory Block of AP-Induced Spontaneous Activity Delayed Maturation of PN Biophysical Properties

Many aspects of functional maturation of MNTB PNs are coordinated with growth of the CH, likely to reduce the excitability of PNs as the weight of their dominant synaptic input increases substantially^30,32,39^. During the first postnatal week, input resistance (R_in_) declines from near 1 GΩ to ∼250 MΩ, and the resting membrane potential (RMP) hyperpolarizes from ∼-64 to −72 mV^30,32,39^. We investigated effects of TeNT via whole-cell current-clamp recordings at the same ages and from the same cells used for recording sEPSCs (Fig. 3 and Table S3). cMNTB PNs displayed larger R_in_ values, measured as slope resistance around RMP (Fig. 3A, A1, and B), which decreased with age and did not become similar to iMNTB PNs until P9, representing an approximate 5-day maturation delay. After ear canal opening at P14, the R_in_ increased in cMNTB PNs, further suggesting homeostatic mechanisms after ear canal opening to increase excitability. The decrease in membrane time constant in cMNTB PNs followed a similar 5-day maturation delay, but remained larger at P9 and P14 (Fig. 3A2 and C). RMP was on average more depolarized in cMNTB PNs across these ages, but due to large variance in these values was not consistently statistically significant (Fig. 3D). During hyperpolarizing current injections, MNTB PNs exhibit a prominent depolarizing sag due to the activation of hyperpolarization-activated cyclic-nucleotide-gated (HCN) channels^47,48^. Due to differences in R_in_ among PNs, we normalized measurement of depolarizing sag to a peak membrane potential of approximately −90mV, enough to elicit depolarizing sags ≥ 2 mV in most cells (Fig. 3A3 and E). Average values were similar for iMNTB and cMNTB, and did not change across all ages. Overall, these data reveal a delayed developmental trajectory that converges toward control values at P9, but with a 5-day delay^30,32,39^.

**Figure 3.**
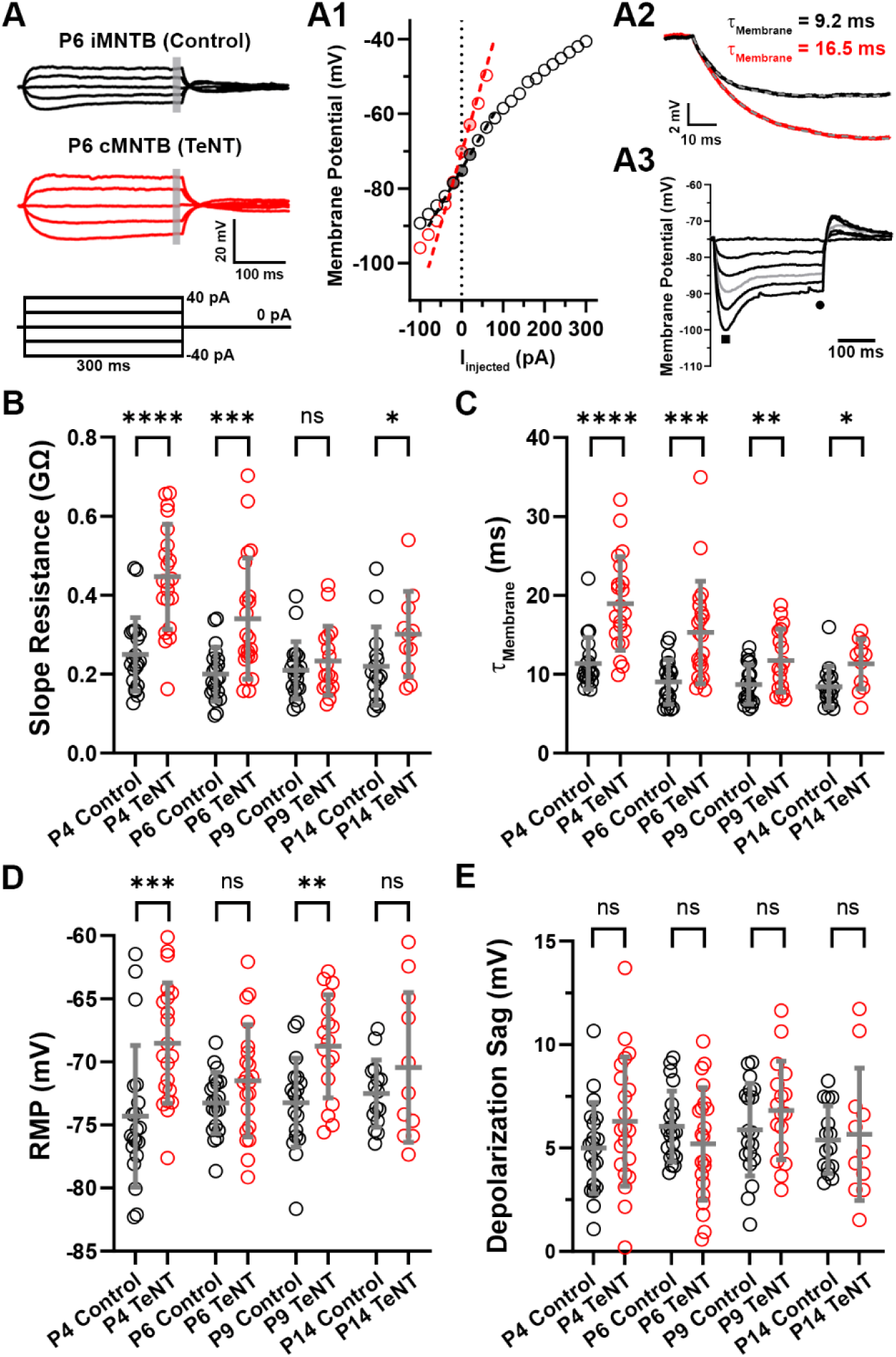
TeNT delays maturation of several physiological properties of MNTB PNs. (A), Exemplary current-clamp traces of P6 PNs recorded from iMNTB (non-transduced, control; black traces and symbols) and cMNTB (transduced, TeNT expression; red traces and symbols). Grey bar indicates steady-state voltages for calculating V-I curve in middle panel. The current command protocol is shown at bottom left. (A1) V-I curves for cells at left. Dashed lines indicate slope resistance fit to three data points surrounding 0 pA. (A2), Exemplary responses to −20 pA hyperpolarization step for P6 iMNTB (black trace) and cMNTB (red trace) PNs to calculate membrane time constant (τ_Membrane_), which was fit by a single exponential (dashed grey line). (A3), Response of P6 iMNTB PN to a series of hyperpolarizing current steps, showing depolarization sag as difference between peak (square) and steady-state (circle) voltages. Values measured for all cells at peak voltage ≈ −90 mV (grey trace). (B-E), Plots showing the slope resistance, membrane time constant, resting membrane potential (RMP), and depolarization sag for iMNTB (Control, black symbols; P4, n = 20; P6, n =20; P9, n = 19; P14, n = 15) and cMNTB (TeNT, red symbols; P4, n = 21; P6, n =21; P9, n = 17; P14, n = 11) PNs across neonatal ages. Mean (horizontal gray lines) and standard deviation (error bars). Significance levels are indicated according to the convention: *P < 0.05, **P < 0.01, ***P < 0.001, ****P < 0.0001, and not significant (ns). Statistical results for panels (B-E) reported in Table S7.

### Pre-Sensory Block of AP-Induced Spontaneous Activity Delayed Maturation of AP Current Thresholds and Slowed AP Kinetics

We next tested if TeNT expression affected AP current and voltage thresholds, and AP waveform kinetics in PNs (Fig. 4 and Table S4). These parameters were quantified from the first suprathreshold spike (rheobase), the threshold current required to elicit a spike, and their phase-plane plots as illustrated by exemplary waveforms at P6 (Fig. 4A1, A2). AP latency was calculated for spikes +40 pA above rheobase to reduce variance, revealing longer values in the cMNTB across postnatal ages (Fig. 4B, C). The threshold current was significantly less in cMNTB than iMNTB PNs from P4-P14, and at P9 was similar to P4 iMNTB, indicating a 5-day maturational delay (Fig. 4D). Voltage thresholds became more hyperpolarized in iMNTB from P4-P14, but did not change in cMNTB from P4-P9, so differences achieved significance at P9 (Fig. 4E). However, hyperpolarization of voltage threshold in cMNTB occurred rapidly after ear canal opening, yielding similar values to iMNTB at P14 (Fig. 4E). AP kinetics were subtly different between iMNTB and cMNTB PNs, because all parameters (amplitude, depolarization rate, and repolarization rate) followed similar developmental increases from P4-P9 (Fig. 4G-I). The net result was that AP half-width was on average longer in cMNTB until P9, although not statistically significant (Fig. 4F). AP kinetics continued to speed up between P9-P14, with amplitudes increasing, and voltage and current thresholds became more similar between groups, likely reflecting experience-dependent adaptation of brainstem neural circuits.

**Figure 4.**
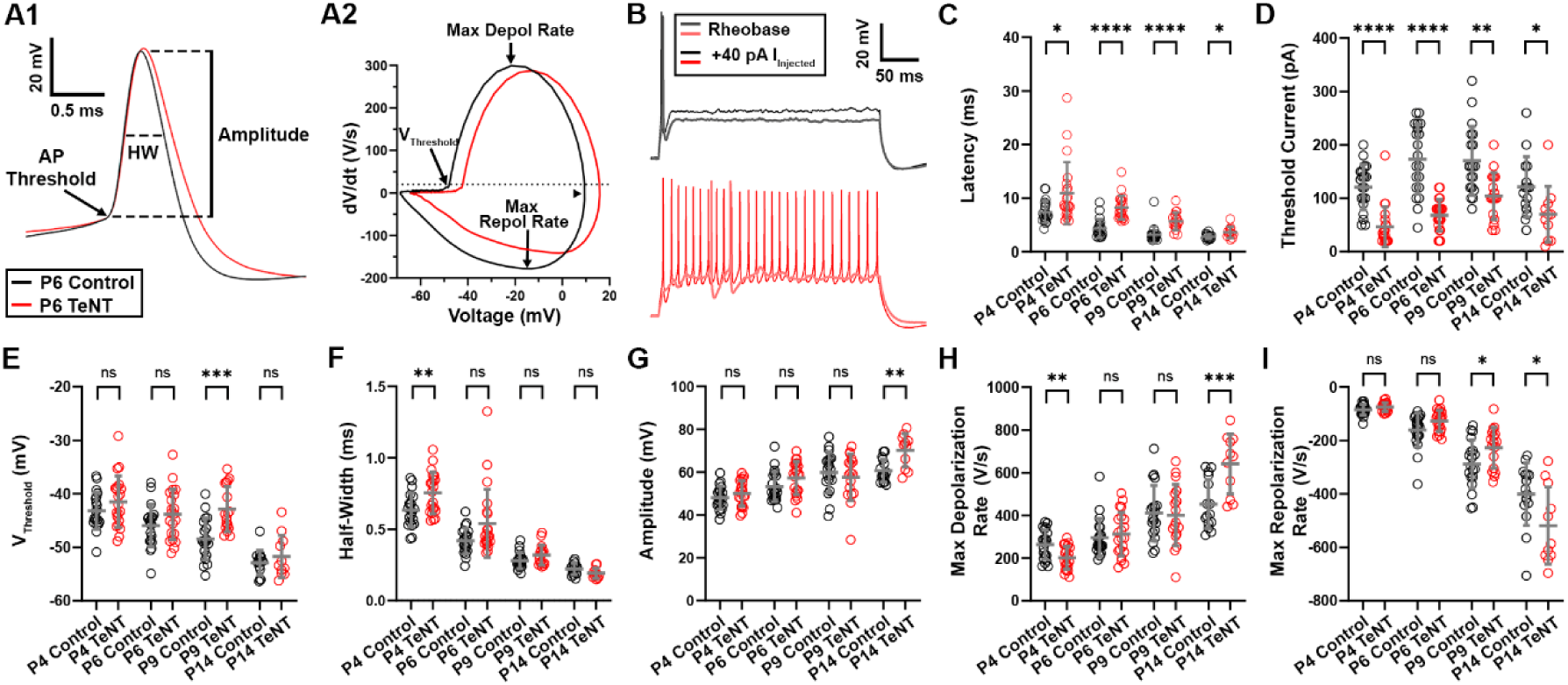
TeNT prevents increase in action potential (AP) current threshold with moderate effects on AP kinetics. (A1) Exemplary action potential (AP) waveforms, aligned at inflection point (AP threshold), recorded at rheobase from P6 PNs in iMNTB (non-transduced control, black trace) and cMNTB (transduced, red trace). AP amplitude and half-width calculations indicated on the traces. (A2) Phase-plane plots for AP waveforms in panel (A). (B), Exemplary current-clamp traces from P6 PNs showing increased spike latency following TeNT expression. Traces are shown at rheobase and at +40 pA injected current step. (C-I), AP latency and current threshold is significantly slower and reduced, respectively in TeNT across ages, and AP kinetic parameters are generally slower but not always significant due to variance in the cell populations (Control, P4, n = 20; P6, n =20; P9, n = 19; P14, n = 15) and cMNTB (TeNT, P4, n = 21; P6, n =21; P9, n = 17; P14, n = 11). Mean (horizontal gray lines) and standard deviation (error bars). Significance levels are indicated according to the convention: *P < 0.05, **P < 0.01, ***P < 0.001, ****P < 0.0001, and not significant (ns). Statistical results for panels (C-I) reported in Table S8.

### Pre-Sensory Block of AP-Induced Spontaneous Activity Prevented Conversion from Tonic to Phasic Spike Pattern

At P3 MNTB PNs begin to transition from a tonic to phasic firing pattern^30,32^ in response to depolarizing current injections. Exemplary responses to hyperpolarizing and depolarizing current injections from P6 iMNTB PNs show the characteristic phasic firing pattern (iMNTB) found by this age in normally developing animals^30,32,49–51^, but tonic firing (cMNTB) following TeNT expression (Fig. 5A, B). For a more quantitative classification of responses, we plotted the number of evoked APs as a function of the injected current. Since many cells entered into depolarization block of APs at large depolarizing currents and thus had non-monotonic profiles, for clarity we plotted values up to the peak response (Fig. 5C). The firing rate at 200 pA best differentiated groups of rate-level curves, and histograms of responses were used to classify MNTB PNs as phasic (≤ 3 APs) or tonic (≥4 APs) (Fig. 5B and C insets; first bin 0-3 spikes). All cMNTB PNs, except three cells, at P4-P14 displayed a characteristic tonic firing pattern (Fig. 5A, B), indicating a failure to transition their firing phenotype. After opening of the ear canals, about 40% of iMNTB PNs at P14 showed a tonic firing pattern, likely indicating, as noted for AP properties, experience-dependent adaptation of brainstem circuitry.

**Figure 5.**
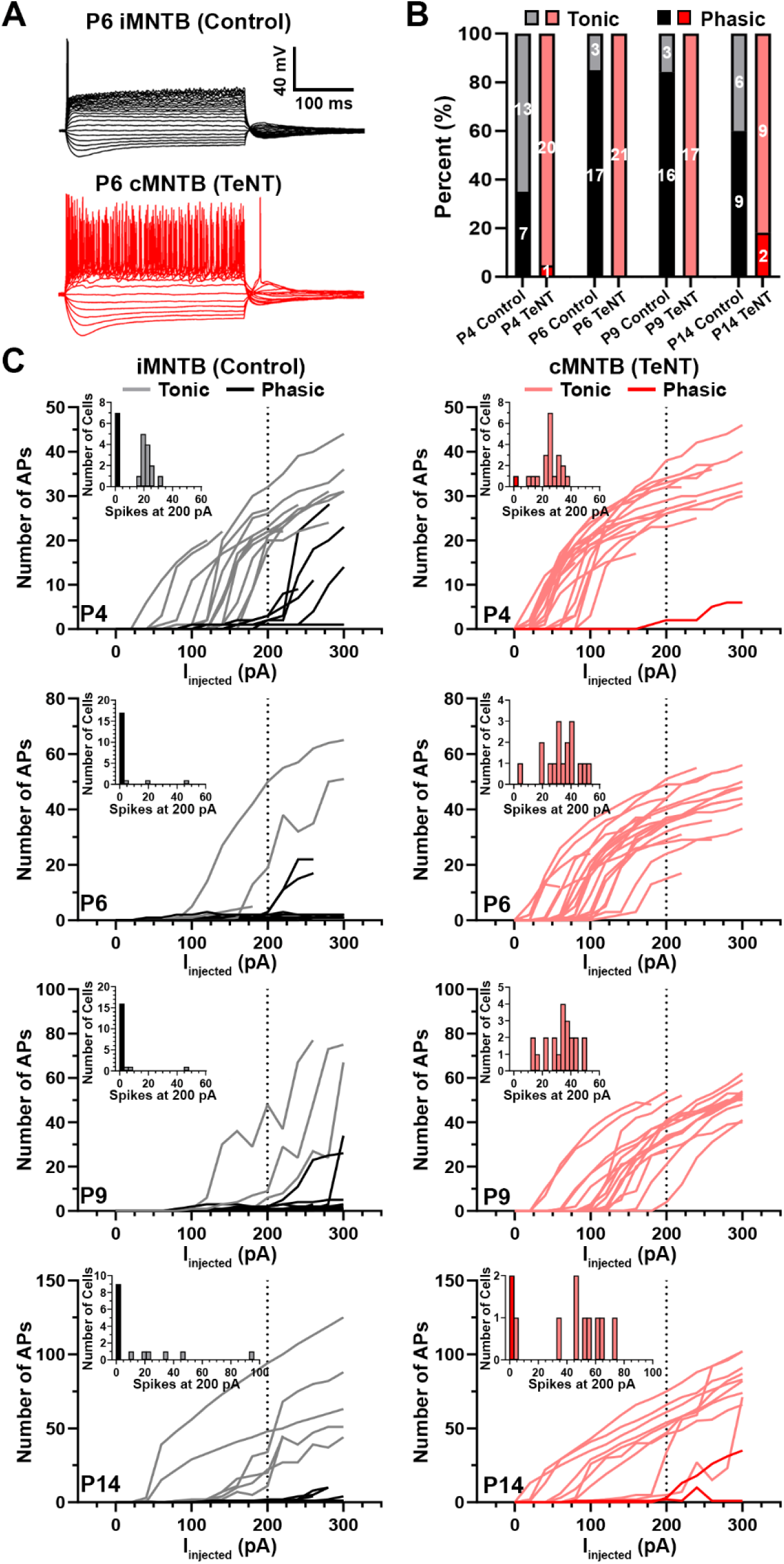
TeNT prevents transition of PNs from tonic to phasic firing at pre-sensory ages. (A), Exemplary current-clamp traces of P6 PNs recorded from iMNTB (non-transduced, control, black traces) and cMNTB (transduced, TeNT; red traces) PNs in response to 20pA current steps from −100 to ≥ 200 pA. (B), Plot showing the percentage of tonic and phasic cells in iMNTB (grey and black) and cMNTB (pink and red). Numbers of cells indicated inside bars. (C), Rate (number of APs) vs injected current across developmental age, utilizing 300 ms depolarizing current steps. Each trace is an individual neuron. For clarity, cells with non-monotonic profiles, likely due to depolarization block, are plotted only until reaching peak spike number. The inset shows a histogram (bin size = 3) of the evoked spike number elicited by a 200 pA current injection. For non-monotonic cells, the maximum number of spikes was used for the histogram. Insets and vertical dotted lines on each plot show utility, especially at pre-sensory ages, in differentiating PNs at ≤ 200 pA current injection as phasic (≤ 3 APs, black or red) or tonic (> 3 APs, grey or pink).

### Pre-Sensory Block of AP-Induced Spontaneous Activity Induced Distinct Physiological Maturational Profiles in the Postsynaptic Neuron

Despite delayed maturation of multiple physiological and biophysical parameters by cMNTB PNs, they converge toward control values by P9 yet maintain a tonic firing phenotype through P14. These data suggest that the lack of neurotransmission from their dominant synaptic input evokes a distinct maturational trajectory that persists even after ear canal opening. To better understand the effects of TeNT expression, we performed a principal component analysis (PCA) of the eleven electrophysiological parameters measured across ages and experimental groups. We held out the phasic/tonic spike rate metric in attempt to understand which combinations of the other parameters could yield the phasic/tonic phenotypes. We anchored this analysis to P9 iMNTB PNs (gold symbols in Figs. 6A, S3B), which represent the endpoint of pre-sensory maturation since ear canals begin to open at P10. The first two principal components (PC1, PC2) accounted for 65.1% of variance among these parameters and reflected general alignment of PC1 with AP parameters and PC2 with passive biophysical properties (see loading vectors in Fig. 6A, B (inset)) and parameter correlations (Fig. S3A). To chart the developmental profile for control PNs, we projected values from P4 and P6 iMNTB PNs electrophysiological parameters onto the axes determined by the P9 iMNTB PNs PCA space (Fig. 6A). These data reveal trajectories upward and leftward from P4-P6, and downward and leftward from P6-P9, with some overlap between ages. K-means clustering analysis was optimal for three clusters, allowing for an unbiased approach to identify maturational groups and account for differences in the rate of maturation among neurons across age (Fig. S3B). Violin plots of the 11 electrophysiological parameters analyzed in the PCA (Fig. S3C) show the maturational changes of iMNTB PNs more smoothly than in age-based plots (Figs. 3, 4), with the most significant changes in biophysical properties from P4-6 and AP properties from P6-P9 (Fig. S3B and C).

**Figure 6.**
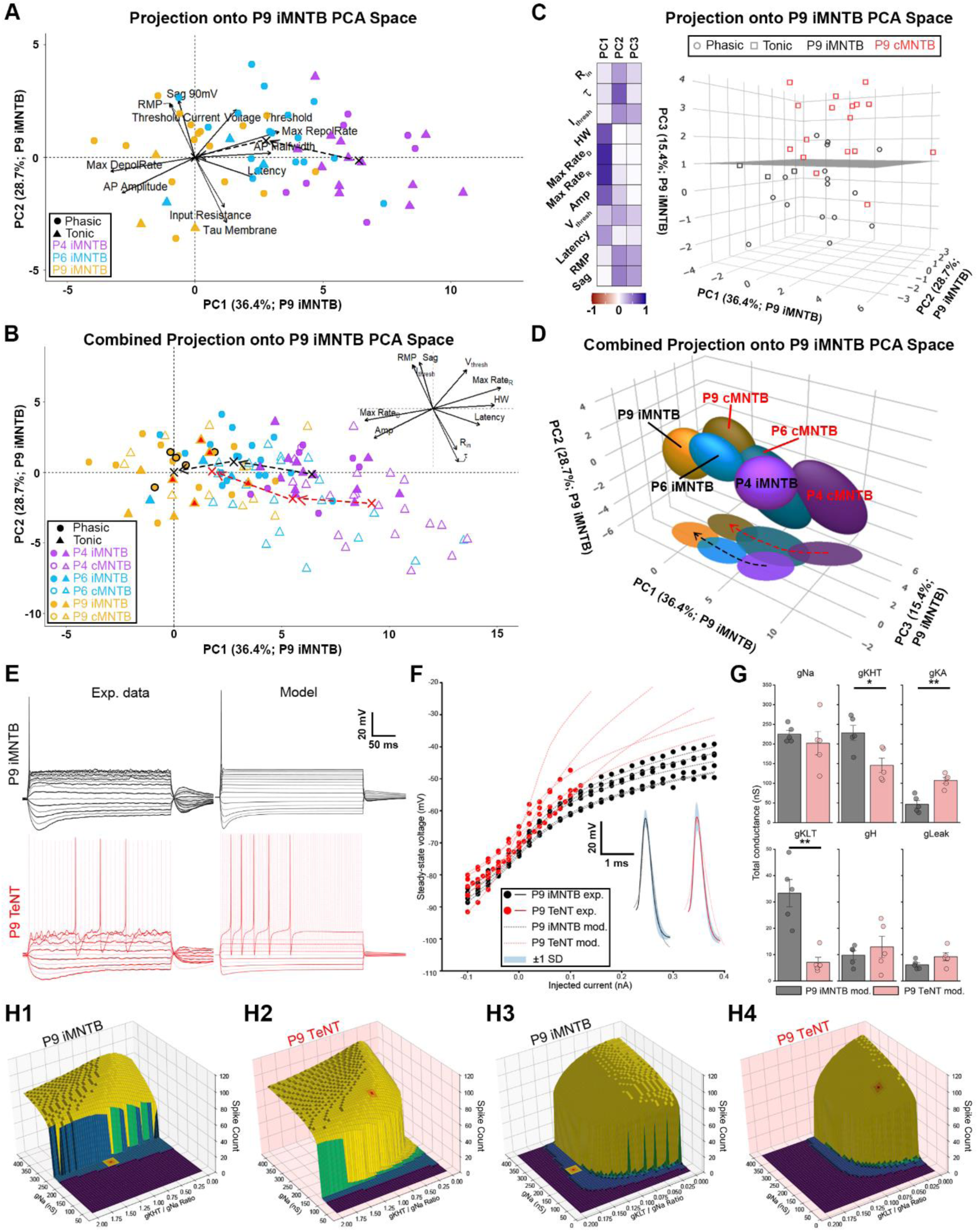
Principal component analysis (PCA) of electrophysiological parameters comparing iMNTB (control) and cMNTB (TeNT expression) PNs, and interpretation of differences using computational modeling. (A), Plot of first two principal components from analysis of P9 iMNTB PNs (n = 19; 11 features). Loading vector for each feature shown with black arrows; length of each arrow corresponds to vector strength. P4 and P6 iMNTB PNs (P4, n = 20; P6, n =20) were projected onto the 2-dimensional PCA space of P9 iMNTB PNs and show maturational trajectory upward and leftward across age (dotted lines; centroids for each age denoted by black “X” symbols). See Methods. Circles and triangles represent phasic and tonic firing patterns, respectively. (B), Values from individual PCA analysis of cMNTB PNs (P4, n = 21; P6, n = 21; P9, n = 17) projected onto 2-dimensional space of P9 iMNTB PNs, and plotted with data from panel (A). Maturational trajectory (dashed red lines; centroids for each age denoted by red “X” symbols) also upward and to left but shifted from iMNTB. Loading vectors from panel (A) shown in inset. P9 iMNTB cells outlined in black and cMNTB cells with red filled symbols used in simulations (panels E-H4). (C), Plot of first three principal components reveals little overlap (compare above and below gray plane) between control and TeNT PNs at P9. Heat map at left shows contribution of variability of each feature for first three PCs. Color and intensity represent degree of correlation. (D), Distinct maturational trajectories in 3D PCA space for iMNTB and cMNTB PNs evident by P4, by visualizing ellipsoids centered and scaled (one standard deviation) for each experimental group, and also projected onto PC1-PC3 plane. (E), Single compartmental (soma) model simulating control (iMNTB; black traces) and TeNT effects in cMNTB (red traces) from P9 PNs. Experimental current-clamp recordings (left column) and computer simulations (model; right column) of representative cells show responses to current steps in 20 pA increments from −100 pA to +200 pA, highlighting traces one step above rheobase. (F), V-I plot of experimental data (filled circles) and simulation (dotted lines) resulting from a Hodgkin-Huxley fit of the passive conductance values (*g^-^*_*H*_, *g^-^*_*KLT*_, and *g^-^*_*Leak*_) for five representative cells from iMNTB (black) and cMNTB (red). Inset: fits for APs at rheobase of active conductance values (*g^-^*_*Na*_, *g^-^*_*KHT*_, and *g^-^*_*KA*_). See Methods. (G), Average values show significant differences in K^+^ conductances between iMNTB and cMNTB. Mean (bars) and standard deviation (error bars). Significance levels are indicated according to the convention: *P < 0.05, **P < 0.01, ***P < 0.001, ****P < 0.0001, and not significant (ns). Mann-Whitney U test (P9 Control vs. P9 TeNT): *g^-^*_*Na*_, P = 0.55; *g^-^*_*KHT*_, P = 3.2E-2; *g^-^*_*KA*_, P = 7.9E-3; *g^-^*_*KLT*_, P = 7.9E-3; *g^-^*_*H*_, P = 0.84; *g^-^*_*Leak*_, P = 0.15. (H1-H4), 3D plots showing the results of 2500 simulations varying *g^-^*_*Na*_, *g^-^*_*KHT*_, *g^-^*_*KLT*_ conductances in cells from panel E. The colors denote the number of Aps (purple: silent; blue: 1 spike; green: 2-3 spikes; yellow: ≥4 spikes). The red square denotes the spatial domain for each simulated PN and the adjacent orange and yellow squares represent deviations with realistic conductance and conductance ratio values, revealing the stability of firing pattern within a large neighborhood of values.

To compare the maturational trajectories of PN physiological properties following TeNT expression, we projected all ages and groups onto P9 iMNTB PCA space (Fig. 6B; iMNTB: closed symbols and cMNTB: open symbols). Interestingly, ∼50% of P4 and P6 cMNTB PNs occupy distinct spatial domains from their iMNTB counterparts, and P9 cMNTB incompletely overlapped P9 iMNTB cells. Given the relatively large variance captured by PC3 (15.4%), we first compared iMNTB and cMNTB at P9, the oldest pre-sensory age, in 3D PCA space (Fig. 6C). Projecting cMNTB onto iMNTB 3D PCA space reveals nearly complete separation between iMNTB and cMNTB PNs (only four iMNTB cells among the main group of cMNTB cells and two cMNTB cells among the main group of iMNTB cells; Fig. 6C). Next, and to simplify the 3D plot of all ages and conditions, each experimental group was represented as an ellipsoid with each ellipsoid dimension one standard deviation from the centroid (Fig. 6D). Ellipsoids for each age and condition (iMNTB, cMNTB) are distinct in space, with significant differences between the Euclidian distances computed from the centroids of all groups (Table S11). The projection of each ellipsoid onto the PC1/PC3 plane is useful to observe that maturational trajectories for iMNTB and cMNTB PNs have diverged by P4 (Fig. 6D). We next asked if P4 cMNTB PNs were similar to younger control ages, since functional maturation of PNs has been charted from as young as E17^30,39^. P4 cMNTB ellipsoids did not overlap ellipsoids of non-injected P0, P1, P2, and P3 PNs, revealing an early divergence in maturational profile after blockade of psSA (Fig. S4A, B). These data indicate that a lack of synaptic input rapidly engenders a cellular program that realigns electrophysiological features to create a compensatory firing phenotype in cMNTB PNs.

### Computational Modeling Predicts that Differential Expression of Kv Channels Underlie Maturational Changes Induced by Pre-Sensory Block of AP-Induced Spontaneous Activity

Given the distinct maturational profiles in PCA space following TeNT expression in CHs, we explored potential ionic mechanisms to explain these events. To simplify this analysis, we created a single-compartment (soma) Hodgkin-Huxley type model incorporating prominent ionic conductances and their parameters that have consensus in the literature and have been utilized in other models of PNs or VCN bushy cells^32,51–55^. These are: sodium conductance (*g^-^*_*Na*_), leak conductance (*g^-^*_*leak*_), low threshold potassium conductance (*g^-^*_*KLT*_), high threshold potassium conductance (*g^-^*_*KHT*_), A-type potassium conductance (*g^-^*_*KA*_), and hyperpolarization activated conductance (*g^-^*_ℎ_). The A-type potassium conductance was explored for a potential role in the longer latency spiking activity observed in cMNTB PNs (Figs. 4B, C and 6E). We chose 5 PNs each from P9 iMNTB and cMNTB that were closest to the centroid for each group (highlighted in Fig. 6B) to investigate differential expression of ion channels that underlie the effects of TeNT expression.

The experimental and modeled responses to hyperpolarizing and depolarizing current steps are shown for one cell from each group (rheobase and at 200 pA injected current highlighted; Fig. 6E). In the first stage of modeling, steady-state V-I curves for each cell were fit individually using parameters for the subthreshold conductances *g^-^*_*leak*_, *g^-^*_*KLT*_, and *g^-^*_ℎ_ (Fig. 6F). Next, APs were fit individually to optimize the values for *g^-^*_*Na*_, *g^-^*_*KHT*_, and *g^-^*_*KA*_ (Fig. 6F, inset). Although individual cells exhibited different levels of all optimized conductances, the averaged response for each group revealed that the greatest differentiator associated with TeNT expression was the level of K^+^ channel expression (Fig. 6G). Reduction of *g^-^*_*KLT*_ in cMNTB was consistent with experimental block of this conductance in PNs, which yields a tonic firing pattern^30,49,50^. The decrease in *g^-^*_*KHT*_ and concomitant increase in *g^-^*_*KA*_ are consistent with differences in AP kinetics and longer AP latency in cMNTB PNs (Fig. 4B, C and 6E). We next explored the stability of modeled conductances in simulations by measuring the firing rate holding average predicted conductance values constant, except for co-varying values for *g^-^*_*KLT*_ or *g^-^*_*KHT*_ with *g^-^*_*Na*_, and plotting results as 3D surfaces (Fig. 6H1-H4). The average predicted values for *g^-^*_*Na*_, *g^-^*_*KLT*_, and *g^-^*_*KHT*_ are indicated in red on the plots. For co-variation of *g^-^*_*KHT*_ and *g^-^*_*Na*_, within a range of ∼200-275 nS for *g^-^*_*Na*_, *g^-^*_*KHT*_ could vary widely (plotted as *g^-^*_*KHT*_ / *g^-^*_*Na*_ ratio) and not induce a tonic response in iMNTB PN, but an increase in *g^-^*_*Na*_ above this range induced a tonic response regardless of *g^-^*_*KHT*_ values (Fig. 6H1). For cMNTB PNs, the average values yielded a tonic response, and only after *g^-^*_*Na*_ <125 nS could a phasic response result, reflecting a shift in the 3D surface to lower *g^-^*_*Na*_values (Fig. 6H2). In simulations that manipulated only *g^-^*_*Na*_ and *g^-^*_*KLT*_, for iMNTB even large increases in *g^-^*_*Na*_ could not induce a tonic response unless *g^-^*_*KLT*_ was reduced (Fig. 6H3-H4). For cMNTB PNs, the simulation could shift from a tonic to phasic response at lower values for *g^-^*_*KLT*_, reflected in a shift of the 3D profile to lower *g^-^*_*KLT*_ / *g^-^*_*Na*_ ratios. These results indicate the ability for *g^-^*_*KLT*_ relative to *g^-^*_*Na*_ to control spike rate over a broad range of *g^-^*_*Na*_ values, and the stability of modeled values for conductances to generate measured phasic or tonic responses. Analysis of *g^-^*_*KA*_ yielded similar results to *g^-^*_*KHT*_ in that large changes in the *g^-^*_*KA*_ / *g^-^*_*Na*_ ratio did not alter the phasic (iMNTB) or tonic (cMNTB) response (data not shown).

### Pre-Sensory Block of AP-Induced Spontaneous Activity Delayed and Impaired Growth of the CH

Given that CH growth is delayed, yet postsynaptic functional maturation occurs but is delayed and altered, we assayed the size and morphology of CHs across developmental age using high resolution confocal microscopy (Fig. 7 and Tables S1, S5). Previous use of TeNT has described presynaptic effects yielding enlarged nerve terminal size and irregular morphology resulting from an increase in synaptic vesicles^24,25,27^. Transduced CHs were visualized by endogenous mCherry expression with antibody amplification, and Vglut1/2 and Map2 antibodies were used to co-label transduced (Vglut1/2) and visualize non-transduced CHs and PNs, respectively (Fig. 7A, B). Low magnification images of the iMNTB and cMNTB at P9 show expression of mCherry mostly restricted to the CHs in the cMNTB (Fig. 7A-B; arrowheads indicate cells with CHs that appear on nearby sections). Quantification of mCherry and Vglut1/2 co-labeling in the cMNTB showed mostly smaller terminals at P4 (punctate labeling, 78% of PNs). Most PNs were innervated by a large calyx-like terminal at P6 (82%) and P9 (92%), nearly all of which co-labeled with mCherry (Table S1), consistent with values achieved by P4 in iMNTB and in control mice^16,56^. After ear canal opening at P14 a larger percentage of non-transduced calyces (∼29%) was present in the cMNTB suggesting possible ectopic innervation^57–59^. In the iMNTB and midline, mCherry expression is localized to the trapezoid body fiber fascicles (ventral acoustic stria) passing through and ventral to the MNTB *en route* to the cMNTB (Figs. 1C, S1), and other superior olive cell and brainstem cell groups^14,60,61^. The high density of labeled fibers, high percentage of PNs contacted by CHs, and high percentage of CHs that are transduced at P6 and P9, along with determination at P6 that only 14±4% (n = 3 animals) of PNs are poly-innervated, indicate resolution of circuit topography to mature values despite expression of TeNT (Figs. 1C-C2, S1, and Table S1). Transduced CHs at all ages (P4-P14) exhibited increased thickness, compared to non-transduced CHs in the iMNTB (Fig. 7C-K). All labeled CHs and their postsynaptic PNs in tissue sections containing both iMNTB and cMNTB were segmented in 3D using virtual reality software (syGlass) to compare their structural features. Non-transduced and transduced CHs had similar volume at P4, but transduced CHs grew little and had less volume than non-transduced CHs at all older ages (P6, P9, P14; Fig. 7L). The postsynaptic PN volume was slightly smaller in the cMNTB compared to the iMNTB across all pre-sensory ages, except P9 (Fig. 7M), with a reduced surface area (P4: 726 μm vs. 784 μm (7.7%); P6: 759 μm vs. 790 μm (4.0%); P9: 791 μm vs. 803 μm (1.5%); P14: 854 μm vs. 875 μm (2.5%)). Because these differences were small and only significant at P4 and P6, they were not incorporated into computational models for P9 PNs. These data indicate the importance of synaptic transmission to support cellular programs for proper structural growth of synaptic partners.

**Figure 7.**
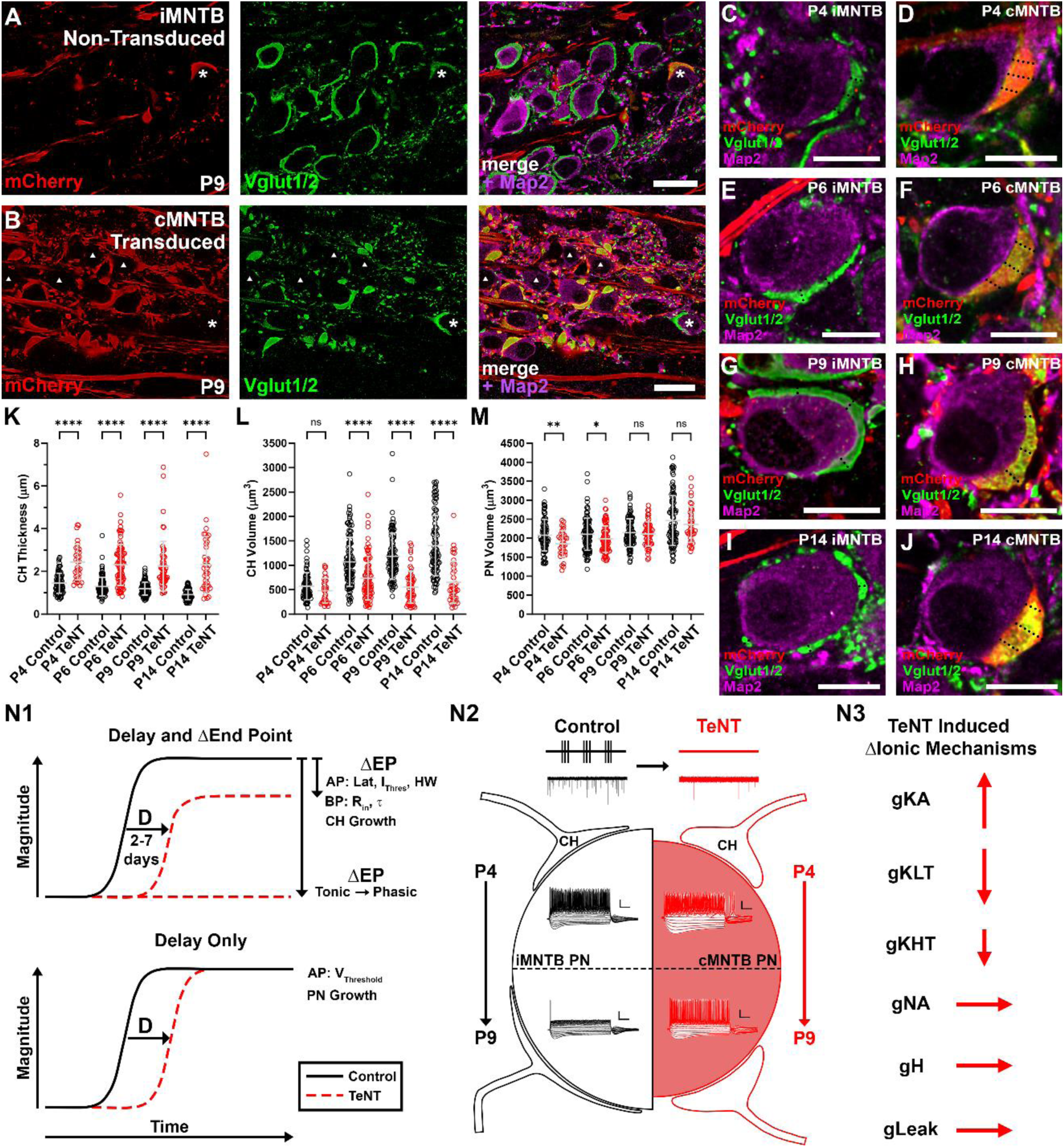
TeNT alters CH and PN morphology. (A-B), Confocal images show transduced (mCherry) CHs in cMNTB and few in iMNTB (asterisk) at P9. Vglut1/2 (green) immunolabeling of CH shows more punctate labeling and thicker calyces in the cMNTB and few non-transduced CHs (asterisk). MNTB PNs immunolabeled with Map2 (magenta). Arrowheads denote cells in cMNTB with a CH outside of image plane. Scale bar: 20 μm. (C-J), Representative 63X confocal images show increased CH thickness in cMNTB and less extension over PN surface across postnatal ages. Dashed lines indicate locations of three thickness measurements averaged from each CH. Scale bar: 10 μm. (K-M), Plots of thickness and volume of non-transduced (control) and transduced (TeNT expression) calyces and MNTB PN volume. Error bars represent the mean ± standard deviation. Significance levels are indicated according to the convention: *P < 0.05, **P < 0.01, ***P < 0.001, ****P < 0.0001, and not significant (ns). Statistical results for panels (K-M) reported in Table S9. (N1-N3), Summary figure showing the effects of TeNT on maturation of structural and biophysical/properties of the CH:PN connection. (N1), Properties delayed with altered endpoints are: anatomical (CH growth); biophysical/physiological (input resistance, membrane time constant, AP latency, AP current threshold, and AP half-width). Tonic to phasic transition does not occur by the oldest pre-sensory age (P9). Properties with delayed but non-altered endpoints are: anatomical (PN growth); biophysical/physiological (AP voltage threshold). (N2), Key effects of TeNT shown at top in abolishing evoked synaptic transmission and reducing spontaneous vesicle release. Cartoon at bottom compares P4 (top half) and P9 (bottom half) showing delayed PN growth, delayed and altered CH growth (smaller and thicker) and failure to transition to phasic firing. (N3), TeNT induced differential ion channel expression, based on computational models, showing increased (*g^-^*_*KA*_) and decreased conductance (*g^-^*_*KLT*_ and *g^-^*_*KHT*_), or no change (*g^-^*_*Na*_, *g^-^*_*H*_, and *g^-^*_*Leak*_). Length of arrow corresponds to the magnitude of change.

## Discussion

In order to study the effects on maturation of synaptic partners by abolishing peripherally-evoked psSA, we developed a viral vector that rapidly-expresses TeNT, at a specific synaptic station along a central sensory pathway, prior to growth and topographic refinement of its driving synaptic input. We utilized the well-defined second to third order connection along the auditory pathway, where the CH grows and mono-innervates PNs in the postsynaptic cell group prior to ear canal opening^14,16–18^. We showed that TeNT expression abolished AP-evoked neurotransmission and resulted in a large reduction in the frequency of sEPSCs, while retaining the ability of the afferent axon to generate spikes (Figs. 1E3, 2B, and S2B-D). Capitalizing on previous characterization of morphological and physiological maturation of the CH:PN connection at daily resolution^16,30,32,39^, we specified delays in maturation of 2-7 days across structural and functional parameters, and in many cases altered maturational endpoints, which in composite caused a failure of PNs to transition from tonic to phasic firing (Fig. 7N1-N2). PCA analysis of PN functional parameters showed that blocking psSA moved PNs into a distinct maturational trajectory evident as early as P4 (Figs. 6B, 6D, and S4). Computational modeling indicates that alterations in expression of K^+^ conductances underlie these experimental observations. We conclude that psSA advances the age at which coordinated circuit maturation begins, yet non-synaptic signals can drive maturation but are insufficient to achieve the normal maturational endpoint.

Our primary endpoint was P9, which is the last age before ear canal opening^18^, when many key CH structural and PN biophysical parameters have plateaued^15,16,30,32,39^ (this study). In well-studied sensory systems such as vision and whisker somatosensation, pre-sensory topographic refinement in central cell groups relies on correlated firing of afferents and Hebbian mechanisms of synaptic strengthening and pruning^6,62^. TeNT expression in olfactory sensory neurons (OSN) from before birth, however, does not alter convergence of OSN axons to appropriate glomeruli, even when target cell types are independently removed, indicating non-Hebbian mechanisms to establish topography^9^. SA patterns instead instruct expression of guidance molecules in the OSN axon^63^. Likewise, TeNT expression in retinal ON-bipolar cells prior to eye opening does not alter their selective innervation of ON-retinal ganglion cells (RGCs)^24^. The formation of CHs, which monoinnervate their target PNs, persists following TeNT expression but with a 2-day delay, indicating that non-Hebbian mechanisms can operate in this system to sculpt circuit topography. Mechanisms such as axonal-sorting molecules^35^ and nerve terminal growth cues^64,65^ may serve as a redundant mechanism to synaptic transmission. psSA, then, is not necessary for CH growth, unless electrical signals in axons, similar to OSNs, were to have effects via undiscovered routes (contact mediated or secreted signaling cues), such as via activation of astrocytes which are intertwined with growing CHs and their axons^66^.

Manipulations of psSA in the auditory system have been aimed at receptor cell function (inner hair cell Otoferlin, Vglut3, Tmc1) and not the primary afferent that enters the CNS^20,67,68^. Notably, genetic manipulation (Vglut3 KO) to block cochlear inner hair cell neurotransmission results in hyperexcitability of auditory nerve fibers through interactions with supporting cells, preserving bursting psSA in the ascending auditory pathway^23^, and suggests that other deafness models may undergo the same homeostatic compensation. The L-type Ca^2+^ channel KO, however, affects inner hair cells and also reduces cell numbers in brainstem cell groups, including PNs in the MNTB^69–71^, but these channels are expressed in many of these same neurons^72,73^ so effects of altered psSA are difficult to isolate. Subtle alteration in the bursting pattern of psSA by manipulating efferent innervation of inner hair cells also did not affect CH formation^74^ or synaptic transmission^75^. Future studies could examine the role of bursting activity patterns, using the detailed, near daily template we have established for CH:PN circuit maturation.

Structural analysis of our secondary endpoint at P14, after ear canal opening, showed a reduction in the number and percentage of PNs in the cMNTB contacted by transduced CHs (Table S1). This observation could reflect CH withdrawal, similar to TeNT-expressing OSNs which extend, several days after initial appropriate innervation, into nearby glomeruli^9^. The increase in non-transduced CHs could also reflect branching of the few initially non-transduced CHs or of ipsilateral axons en route to the cMNTB, as occurs following manipulations that lesion GBCs unilaterally^57,59^. Interestingly, when TeNT was expressed in CHs after their formation with likely post-hearing onset, it did not result in changes in CH structure^26^. TeNT expression was driven in that study by the *Math5* (*Atoh7*) promoter^76^, which may have resulted in a bias toward transducing SBCs relative to CH-forming GBCs^77^, contributing to their observations of normal CH structure.

Capitalizing on the homogeneous composition of PNs in the MNTB of mice^78,79^ and performing *ex vivo* recordings across multiple developmental time points, we propose specific effects of psSA in developmental regulation of K_V_ channel expression. In organotypic brain slice, albeit from older (P9-P12) animals, CHs degenerate, which creates a situation similar to TeNT expression of reduced activity from the dominant excitatory input. PNs then exhibit decreased K_LT_ and K_HT_ expression and acquire a tonic phenotype^80^ that is prevented by Ca^2+^ signaling via tonic depolarization in high K^+^. K_HT_-regulation occurs via a CREB phosphorylation-based mechanism typically linked to experience-dependent plasticity^81–83^. One question in our study is why functional maturation of the PN is delayed and altered rather than frozen by TeNT at approximately the P2 time point of its earliest expression. One possibility is that the TeNT-induced delayed decrease in R_in_, mediated by a reduction in K_LT_ expression, may permit even the residual sEPSPs to drive depolarization sufficient for postsynaptic Ca^2+^ signaling through L-type voltage gated Ca^2+^ channels^73^ or Ca^2+^ permeable AMPA receptors^84^. A possible slow accumulation of Ca^2+^-based molecular signals could result in a maturational delay. Alternatively, non-synaptic signals that foster CH growth may activate PN intrinsic cellular programs, or PNs may detect the increasing physical contact area of the growing CH. As a consequence of reduced K_LT_, a concomitant increase in the A-type potassium conductance was observed in our computational modeling, which partially preserved the developmental decrease in excitability (Figs. 3B, 6G)^30,32,39^. Previous electrophysiological studies in mice have shown the presence of K_A_ conductance in PNs, mediated by K_V_4 subunits^85^ and single-cell sequencing revealed high levels of *Kcnd2* (K_V_4.2) expression in P3 PNs^72^. The kinetics of K_V_4 channels allow them to shape AP waveforms^86^, and in our models they compensated for reduced K_HT_ and increased AP latency in cMNTB PNs. In PNs Kv4 channels are likely expressed in the dendrites^85,87^. Ultrastructural analysis from electron microscopy volumes have shown PNs have highly branched dendritic arbors at P2, which are pruned back by P9^16^ (Unpublished data). Thus, TeNT expression may prevent dendrite pruning and elevate Kv4 expression compared to iMNTB PNs. In summary, this study isolated effects of psSA, which provide a foundation to further investigate neurotransmission and cell contact-based signaling along with high-resolution structural analysis at the single neuron level to further dissect maturation programs for ion channel expression and structural dynamics that define neural circuit formation.

## Methods

### Animals

All procedures involving animals were approved by the University of South Florida Institutional Animal Care and Use Committee. FVB mice (NCI; Frederick, MD and Jackson Laboratory; Bar Harbor, ME; RRID:IMSR_ARC:FVB) of either sex were used in all experiments.

### DNA Construct and Recombinant Viral Vector Production

A recombinant Helper-dependent Adenovirus (HdAd28E4) plasmid was optimized for rapid, high-level expression of the 50 kDa tetanus neurotoxin light chain (TeNTLC), a metalloprotease that cleaves the synaptic vesicle fusion protein synaptobrevin-2^29^, independently of the fluorescent reporter mCherry. The dual expression plasmid utilizes the pUNISHER cassette^28^ to drive expression of TeNTLC along with a separate neuron specific mCherry expression cassette driven by the 470 bp hSyn promoter. Production of the HdAd was carried out as previously described^88^. For stereotaxic viral injections the amount of the virus injected did not exceed a total of 2.03×10^8^ viral particles/μL. Preparation of the viral injection solution followed the protocol previously described^89^, with the virus stock solution diluted in storage buffer (250 mM Sucrose, 10 mM HEPES, 1 mM MgCl_2_ dissolved in nanopure H_2_O) and 20% mannitol solution.

### Stereotaxic Viral Injection

Viral vector injections were performed on newborn (postnatal day (P)0) pups as previously described^89^. Anesthesia was induced by cryoanesthesia (deep hypothermia). The depth of anesthesia was verified by toe pinch and a lack of ensuing reflex response. Pups were positioned on a chilled, clay-filled aluminum block (custom built) in the prone position and held in place with additional clay. The aluminum block was positioned on a Kopf stereotaxic frame (model 940 digital; David Kopf Instruments, Tujunga, CA). A small incision was made in the skin at the injection site using a sterile 26G needle. Micropipette glass capillary needles (Blaubland; IntraMARK) were pulled using a Narishige puller (model PC 10, RRID:SCR_022057), clipped to an approximate tip diameter of 20-50 µm, and loaded with the virus solution (HdAd28E4 Cpun TeNTLC syn mCherry) using capillary action. For P0 mouse pups lambda and bregma (where coronal and sagittal sutures intersect) were identified visually by evident suture lines in the skull. An injection glass needle was positioned directly above lambda and slowly moved to bregma assuring proper anterior/posterior stereotaxic alignment of the mouse pup. From bregma (location considered 0.0 mm) the coordinates for the injected ventral cochlear nucleus are as follows: anterior-posterior: −5.1 mm, medial-lateral: −1.4 mm, and dorsal-ventral: −2.9-3.2 mm (assuming a 3.5mm distance from lambda to bregma) (Fig. 3.3A). The virus solution (up to 1 μL, but typically 200-500 nL) was then slowly infused at a rate of 0.5 μL/min. After injection of virus, the glass capillary needle was left in place for 1 minute prior to slow withdrawal. The animal was placed under a heat lamp (∼35°C) to recover and restore physiological temperature. Pups were rolled in bedding material to ensure that they were accepted by their mother upon return to the cage.

### Slice Preparation for Electrophysiology Experiments

Acute brainstem slices were prepared from neonatal mice as previously described^15,30^. FVB mouse pups (P0-P14) were decapitated, and the brain was immediately dissected in ice-cold, low Ca^2+^ artificial cerebrospinal fluid (ACSF) containing the following (in mM): 125 NaCl, 2.5 KCl, 3 MgCl_2_, 0.1 CaCl_2_, 25 glucose, 25 NaHCO_3_, 1.25 NaH_2_PO_4_, 0.4 ascorbic acid, 3 myo-inositol, and 2 sodium pyruvate, pH 7.3. Coronal 250-300 μm brainstem slices containing the MNTB were cut using a vibratome (VT1200S, Leica), stored at 37°C for 1 hr and kept at room temperature (RT) in normal ACSF containing the following (in mM): 125 NaCl, 2.5 KCl, 1 MgCl_2_, 2 CaCl_2_, 25 glucose, 25 NaHCO_3_, 1.25 NaH_2_PO_4_, 0.4 ascorbic acid, 3 myo-inositol, and 2 sodium pyruvate, pH 7.3 until experimentation. All solutions were constantly bubbled with 95% O_2_/5% CO_2_.

### Electrophysiology

For whole-cell recordings (P4-P14) following viral vector injection, 7-12 animals per age from at least 5 litters were used for data analysis. All electrophysiology recordings were performed at near physiological temperature (34-36°C). Solution temperature was regulated using a dual-channel temperature controller with both in-line and chamber heaters (TC-344B, Warner Instruments). ACSF was constantly bubbled with 95% O_2_/5% CO_2_ and perfused over the slice at a rate of ∼2 mL/min using a peristaltic pump (P720, Instech). Neurons were visualized and targeted for whole-cell recordings using a two-photon microscope (HyperScope, Scientifica) equipped with Dodt Gradient Contrast (DGC) optics and an EM-CCD or sCMOS camera (C9100 or ORCA Fusion-BT, respectively; Hamamatsu). All recordings were made from the medial 1/3, high frequency region of the MNTB to minimize the effects of differential ion channel expression and maturational gradients along the tonotopic axis^90^. In the cMNTB, neurons postsynaptic to labeled CHs (mCherry co-expressed with TeNT) were targeted for recording, except at P4, when the presence of large labeled fluorescently labeled calyces was less frequent (4/21 recordings). The other recordings (17/21) were made from PNs without an identifiable CH input to achieve a comparable sample size for statistical analysis. Patch-pipettes were pulled to a tip resistance of 2-4 MΩ for postsynaptic recordings (PNs) and 5-7 MΩ for presynaptic recordings (CH) using a micropipette puller (P-1000, Sutter Instrument Co.). The internal recording solution contained the following (in mM): 114 potassium gluconate, 26 KCl, 2 MgCl_2_, 0.1 CaCl_2_, 1.1 EGTA-Na_4_, 10 Hepes, 5 sodium phosphocreatine, and 4 ATP-Mg, pH 7.3. In some recordings the intracellular solution included a 10,000 MW dextran-conjugated fluorophore (Alexa Fluor 488 or 647; 50 μM) for visual identification. Whole-cell voltage and current-clamp recordings were made using a patch-clamp amplifier (model EPC 10 USB, HEKA Electronik, RRID:SCR_018399) and data were acquired at a sampling rate of 50 kHz and low-pass filtered at 3 kHz. For voltage-clamp protocols series resistance during pre- and postsynaptic recordings was in the range of 8-25 MΩ and 4-10 MΩ, respectively and compensated 50-90% with a lag of 10 μs. For current-clamp recordings, pipette capacitance neutralization and bridge balance were adjusted and monitored during all recording sessions. Potentials were corrected online for a −13 mV liquid junction potential, calculated at 35°C using pCLAMP 9.2 software (RRID:SCR_011323)^30^. For all recordings the glycine receptor antagonist Strychnine (2 μM) (European Pharmacopoeia) and GABA_A_ receptor antagonist SR 95531 hydrobromide (gabazine; 10 μM) (Alomone Labs, RRID:SCR_013570) were added to the extracellular solution to block inhibitory glycinergic and GABAergic currents, respectively^91^. Spontaneous excitatory postsynaptic currents (sEPSCs) were recorded without application of the voltage-gated sodium channel blocker tetrodotoxin (TTX), so that action potential waveforms could be analyzed for all cells. Previous reports and our own unpublished results have shown sEPSCs kinetics were unaffected by TTX^39,92^.

### Electrophysiology Data Analysis

Data was analyzed offline with custom routines using Igor Pro (Wavemetrics, RRID:SCR_000325). Individual sEPSC events were detected utilizing a sliding template algorithm^93,94^ with custom code implemented in Igor Pro. The sEPSC template parameters included the peak amplitude, 20-80% rise time, and decay time constant (τ) and were defined based on average values recorded from MNTB PNs at physiological temperature. The detection threshold for individual events was set at 1-2 standard deviations above the baseline noise. Quantification of the rise time and decay time constant (τ) were determined by fitting a single exponential function to individual events. The input resistance of MNTB PNs was calculated around the resting membrane potential by plotting the voltage-current relationship from the steady-state and fitting a regression line to three points. Measurement of action potential (AP) waveform kinetics were made at rheobase, determined from 300 ms current steps at 10-20 pA increments. AP amplitudes were measured and reported from the voltage threshold. The voltage threshold was determined as the membrane potential at which the dV/dt exceeds 10 V/s (P0-P2) and 20 V/s (P3-P14). The AP half-width was determined using the voltage threshold to peak amplitude measurement.

### Principal Component Analysis (PCA)

From the electrophysiological recordings (n = 118 total cells) comparing ipsilateral (control) and contralateral (TeNT expression), to the injection site, MNTB PNs at P4, P6, and P9, a data matrix (118 x 11) was created based on 11 electrophysiological parameters. Additionally, for comparison to non-injected MNTB PNs a 40 x 11 data matrix was created (P0: n = 9 cells; P1: n = 11 cells; P2: n = 10 cells; P3: n = 10 cells) utilizing the same 11 electrophysiological parameters. The 11 parameters used in the analysis were input resistance, membrane time constant, threshold current, AP half-width, AP amplitude, AP voltage threshold, AP latency, AP maximum depolarization rate, AP maximum repolarization rate, resting membrane potential, and depolarization sag. Data were centered and scaled to unit variance (z-score standardization). For projection onto precomputed PCA space (original dataset: P9 iMNTB), the data matrix for individual groups (e.g. P4 iMNTB and P6 iMNTB) was centered and standardized using the mean and standard deviation of the P9 iMNTB dataset and PC scores were computed based on the eigenvalues and eigenvectors of the original dataset PCA space. The optimal number of clusters for the K-means algorithm was determined utilizing the within sum of squares (WSS) method. PCA and K-means clustering were performed in Rstudio (RRID:SCR_000432; R version 4.4.2) utilizing the *FactoMineR* and *factoextra* packages, respectively. For data visualization the *factoextra* and *ggplot2* packages were utilized. Statistical significance of the PCA results between groups was computed using Permutational Multivariate Analysis of Variance Using Distance Matrices (PERMANOVA) implemented using the *vegan* package with Euclidian distances calculated between group centroids. The complete code is available at https://github.com/NikollasBenites/R_analysis_Heller_et_al_2025.

### Computational Neuron Modeling

The MNTB PN model consists of a single compartment (soma) implemented using the NEURON simulation software version 8.2.6 (RRID:SCR_005393)^95^. The Hodgkin-Huxley type model was based on previous numerical simulations of MNTB PNs^32,51,53,54^. The PN model consisted of a single electrical compartment with a membrane capacitance (*C_m_*) connected in parallel with a sodium current (*I_Na_*), leak current (*I_Leak_*), high-threshold potassium current (*I_KHT_*), low-threshold potassium current (*I_KLT_*), A-type potassium current (*I_KA_*), and hyperpolarization-activated cation current (*I_h_*). The changes to the membrane potential (*V*) were described by the following differential equation:

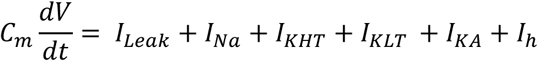

The leakage current was modeled by the following equation:

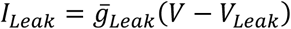

where *g^-^*_*Leak*_ is the maximum steady-state conductance and *V*_*Leak*_ is the reversal potential for the leak current. The voltage dependence of the sodium current was modeled by the following equation:

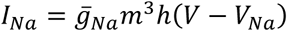

where *g^-^*_*Na*_ is the maximum steady-state conductance, *V*_*Na*_ is the reversal potential for the sodium current, and *m* and *h* are gating variables that model the open probability of the channel. The gating variables (*m* and *h*) are dimensionless parameters that model activation (*m*) and inactivation (*h*) of the sodium channel. For the MNTB PN model the outward potassium current was simulated utilizing a high-threshold K_v_3.1-like current and a low-threshold K_v_1.1 and K_v_1.2-like current, and a K_V_4-like current (A-type). The following equations modeled the voltage dependence of the *I_KHT_*, *I_KLT_*:

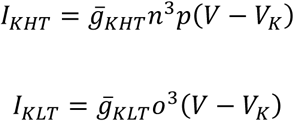

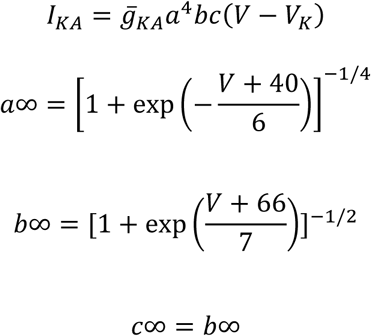

and the *I_KA_*:

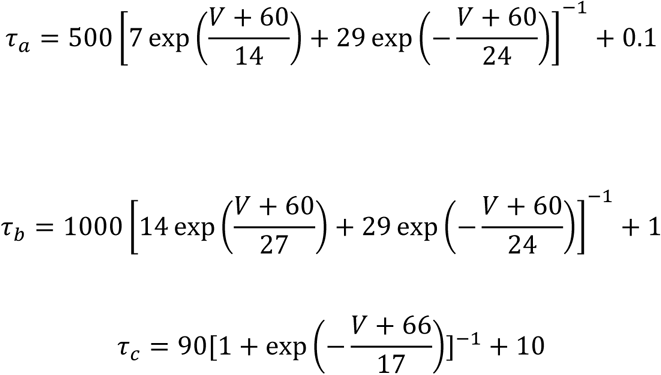

where *g^-^*_*KHT*_, *g^-^*_*KLT*_, and *g^-^*_*KA*_ are the maximum steady-state conductance for the high and low-threshold potassium current, and the A-type current respectively. The gating variables model the activation and inactivation (*n* and *p*) of the high-threshold potassium channel, and activation (*o*) of the low-threshold potassium channel. For the A-type current, the steady-state activation *a∞* and inactivation *b∞*, and *c∞* variables were described using Boltzmann-type equations, modeled as sigmoidal functions of the membrane voltage, with slopes and V1/2 values based on experimental fits adapted from Rothman & Manis (2003). The time constants for each gating variable, *τ_a_*, *τ_b_*, and *τ_c_* were voltage-dependent and derived from empirical fits to experimental data^55^. The hyperpolarization-activated current was modeled by the following equation:

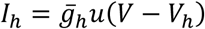

where *g^-^*_ℎ_ is the maximum steady-state conductance for the *I_h_* current, and the gating variable *u* models the activation. The time-dependence for each gating variable was governed by the following differential equation:

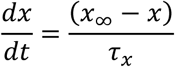

where *x* corresponds to the gating variables for each ion channel modeled (*m, h, n, p, o, a, b, c,* and *u*) and for a given voltage *x* approaches a steady-state value *x*_∞_ with a time constant τ_*x*_. The steady-state open probability value for each ion channel, *x*_∞_, (except *I*_*KA*_) was calculated from the following equation:

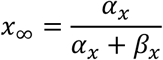

where α_*x*_ and β_*x*_ are the activation and inactivation rate constants, respectively for each ion channel (*x = m, h, n, p, o,* and *u*). For the current through each ion channel, except I_KA_, the voltage dependency of the rate constants was described from the following equations:

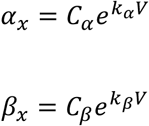

where *C*_α_, *C*_β_ *k*_α_, and *k*_β_ are the constants that determine the rates and voltage dependence (*V*) of current activation. The parameters for voltage activation rate constants for each ion channel are listed in Table S12. The reversal potential for each ion channel modeled was: *E_Leak_* = −73 mV, *E_Na_* = 62.77 mV, *E_KHT_* = −106.81 mV, *E_KLT_* = −106.81 mV, and *E_h_* = −45 mV. To quantitatively match experimental recordings, model parameters were fitted using custom Python scripts coupled to NEURON. The fitting was performed in two stages: (1) for the subthreshold conductances, fitting of the V-I profile at the steady-state membrane potentials, and (2) suprathreshold conductances, fitting of action potential waveforms at the rheobase. For the first stage, *g^-^*_*Leak*_, *g^-^*_*KLT*_, *g^-^*_*H*_, and *E*_*Leak*_ were optimized by minimizing the error between simulated and experimental steady-state voltage responses to the same range of experimental current injections, using the explained sum of squares (ESS):

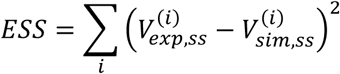

Where *V*^(*i*)^_exp,ss_ is the mean of the experimental voltage during the final 50 ms of the step, *V*^(*i*)^_sim,ss_ is the simulated mean voltage during the same period, and (*i*) is the index over all current injection steps. After fixing the subthreshold parameters, the difference between experimental and simulated voltage traces was quantified using mean squared error (MSE) to optimize the suprathreshold conductances (including *g^-^*_*Na*_, *g^-^*_*KHT*_, and *g^-^*_*KA*_), computed as the average squared pointwise deviation across the full trace. Additional penalty terms were included for mismatches in spike features and for temporal misalignment between AP peaks. The overall cost function was a weighted sum of the trace MSE, feature penalties, and alignment penalties:

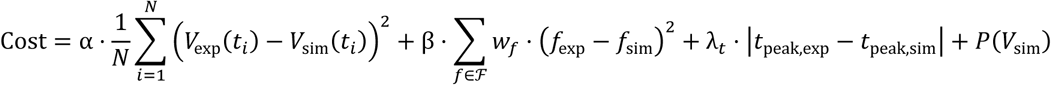

Where *N* is the total number of time points in the voltage trace, and *V*_exp_ and *V*_sim_ represent the experimental and simulated membrane potentials, respectively, at each time point *t*_*i*_. The first term of the cost function quantifies the mean squared error between the full voltage traces scaled by a factor *α*. The second term penalizes mismatches in specific action potential features (e.g., spike threshold, peak voltage, latency) scaled by a factor *β*, and *w*_*f*_ is a tunable weight assigned to each feature *f* ∈ ℱ. The third term accounts for temporal misalignment between the experimental and simulated action potential peaks, scaled by a factor λ_*t*_. The final term, *P*(*V*_sim_), represents a regularization penalty that discourages biologically implausible voltage traces, such as excessive depolarization or spontaneous activity. Optimization was performed using both global (differential evolution) and local (gradient-based) methods via *SciPy*. All simulations were performed at 35°C with a time step of 0.02 ms.

### Immunofluorescence Imaging

FVB mice (P2-P14) were anesthetized using hypothermia/cryoanesthesia (P2) or with Avertin (P4 and older; 250 mg/kg intraperitoneal injection) and perfused transcardially with 10 mM phosphate-buffered saline (PBS), pH 7.4, followed by a solution of 4% paraformaldehyde (PFA) in PBS. Each brain was removed from the skull and post-fixed overnight at 4°C in 4% PFA in PBS and kept in 0.4% PFA until sectioning (up to 1-2 weeks). The brains were transferred to cryoprotectant (30% sucrose in PBS) at 4°C for 1-2 days prior to freezing and sectioning. Coronal sections of the brainstem were cut at ∼50-60 μm thickness using a freezing microtome (model HM 450, Microm, Waltham, MA). Epitope retrieval was performed in 10mM citric acid, pH 6.0, at 95°C for 30 minutes before sections were moved into blocking solution (3% donkey serum in PBS containing 0.1% Triton X-100) for 1 hour. The sections were then incubated overnight at 4°C on an orbital shaker with primary antibodies diluted in 3% donkey serum in PBS. Primary antibodies included: dsRed (1:500, Cat# 632496: Takara Bio, RRID:AB_10013483) to amplify endogenous mCherry signal, microtubule associated protein 2 (Map2, 1:2500, catalog # CPCA-Map2; EnCor Biotechnology) to label MNTB PNs, vesicular glutamate transporter 1 (Vglut1, 1:2500, Cat# AB5905; EMD Millipore) and Vglut2 (1:2500, Cat# AB2251-I) to label the CH. After primary antibody incubation slices were washed three times (5 min each) with PBS followed by incubation with the appropriate secondary antibodies (Molecular Probes, Grand Island, NY; Jackson Immunoresearch Laboratories, West Grove, PA) diluted in blocking solution at 1:500 for 2 h at RT. Slices were washed three times (5 min each) with PBS before being imaged on an inverted confocal microscope equipped with a motorized stage (SP8 LIGHTNING Confocal Microscope, RRID:SCR_018169) using a Leica 63X HC PL APO/1.4 NA oil immersion objective. Z-stacks were collected in 0.3 μm steps. Scanning parameters (e.g. laser power and gain) were optimized based on fluorescence signal intensity from the MNTB contralateral to the injection site (transduced, TeNT expression) and the same parameters were utilized for image acquisition of the ipsilateral MNTB (non-transduced, control). Images were postprocessed with Lightning deconvolution using Leica imaging software (Leica Application Suite X, RRID:SCR_013673) for increased contrast and resolution.

### Segmentation of CHs and PNs from Confocal Image Stacks

Confocal image volumes were imported into the 3D virtual reality software syGlass (www.syglass.io, IstoVisio, Inc; RRID:SCR_017961) for manual segmentation and reconstruction of calyces and MNTB PNs. Mice (n = 2-3 per age) from at least two different litters per age were processed for immunohistochemistry staining and confocal imaging. For each image stack all CHs and associated PN that were fully contained within the confocal image volume were segmented for quantitative analysis. For manual segmentation a region of interest was drawn around each CH:PN. The 3D “painting tool” in syGlass was utilized to manually segment each structure and export surface meshes for quantitative analysis of the volume and surface area for each CH and associated MNTB PN. Thickness measurements of each CH were quantified using the measure tool in ImageJ (RRID:SCR_003070). To quantify the thickness, given the irregular morphology, three measurements were taken through the approximate centroid of the terminal and average values were reported.

### Statistics

All statistical analyses were performed in GraphPad Prism (GraphPad Software, San Diego, CA, United States). The normality of datasets was analyzed using the Shapiro-Wilk test, with parametric or non-parametric statistical tests carried out accordingly. To compare two groups, an unpaired t-test (parametric), Mann-Whitney test (non-parametric), or Wilcoxon signed-rank test (non-parametric) was carried out. To compare three or more groups, one-way ANOVA with Tukey’s test for multiple comparisons or Kruskal-Wallis test with either Dunn’s test or Wilcoxon test for multiple comparisons was carried out for parametric or non-parametric data. All values indicated in the results are presented as the mean ± standard deviation. Significance levels reported in the results and figures are indicated according to the following convention: *P < 0.05, **P < 0.01, ***P < 0.001, ****P < 0.0001, and not significant (ns).

## Acknowledgments

This work was supported by NIH/NIDCD grant R01 DC007695 to GS, HvG, and SY. We thank Dr. Paul Manis for discussion of computational modeling strategies and results. Imaging experiments were performed in the University of South Florida Lisa Muma Weitz Laboratory for Advanced Microscopy & Cell Imaging Core, with advice from Dr. Byeong “Jake” Cha. We acknowledge Flepp Wallace and Bret Nowakowski for assistance in calyceal and principal neuron segmentation.

## Author Contributions

Conceptualization, D.T.H, N.M.B., S.M.Y., and G.A.S.; performed research, D.T.H., N.M.B., and E.M.A.; analyzed and interpreted data, D.T.H., N.M.B., and G.A.S.; software, D.T.H., N.M.B., and A.D.; writing – original draft, D.T.H., N.M.B., and G.A.S.; writing – review & editing, D.T.H., N.M.B., E.M.A., A.D., S.M.Y., H.v.G., and G.A.S.; funding acquisition, G.A.S., H.v.G., and S.M.Y.

## References

1 Pumo, G. M., Kitazawa, T. & Rijli, F. M. Epigenetic and Transcriptional Regulation of Spontaneous and Sensory Activity Dependent Programs During Neuronal Circuit Development. Front Neural Circuits 16, 911023 (2022). 10.3389/fncir.2022.911023

2 Blankenship, A. G. & Feller, M. B. Mechanisms underlying spontaneous patterned activity in developing neural circuits. Nat Rev Neurosci 11, 18–29 (2010). 10.1038/nrn2759

3 Kersbergen, C. J. & Bergles, D. E. Priming central sound processing circuits through induction of spontaneous activity in the cochlea before hearing onset. Trends Neurosci 47, 522–537 (2024). 10.1016/j.tins.2024.04.007

4 Martini, F. J., Guillamon-Vivancos, T., Moreno-Juan, V., Valdeolmillos, M. & Lopez-Bendito, G. Spontaneous activity in developing thalamic and cortical sensory networks. Neuron 109, 2519–2534 (2021). 10.1016/j.neuron.2021.06.026

5 Molnar, Z., Luhmann, H. J. & Kanold, P. O. Transient cortical circuits match spontaneous and sensory-driven activity during development. Science 370 (2020). 10.1126/science.abb2153

6 Anton-Bolanos, N. et al. Prenatal activity from thalamic neurons governs the emergence of functional cortical maps in mice. Science 364, 987–990 (2019). 10.1126/science.aav7617

7 Huberman, A. D. et al. Eye-specific retinogeniculate segregation independent of normal neuronal activity. Science 300, 994–998 (2003). 10.1126/science.1080694

8 Shatz, C. J. & Stryker, M. P. Prenatal tetrodotoxin infusion blocks segregation of retinogeniculate afferents. Science 242, 87–89 (1988). 10.1126/science.3175636

9 Yu, C. R. et al. Spontaneous neural activity is required for the establishment and maintenance of the olfactory sensory map. Neuron 42, 553–566 (2004). 10.1016/s0896-6273(04)00224-7

10 Li, H. et al. Laminar and columnar development of barrel cortex relies on thalamocortical neurotransmission. Neuron 79, 970–986 (2013). 10.1016/j.neuron.2013.06.043

11 Babola, T. A. et al. Purinergic Signaling Controls Spontaneous Activity in the Auditory System throughout Early Development. J Neurosci 41, 594–612 (2021). 10.1523/JNEUROSCI.2178-20.2020

12 Tritsch, N. X. et al. Calcium action potentials in hair cells pattern auditory neuron activity before hearing onset. Nat Neurosci 13, 1050–1052 (2010). 10.1038/nn.2604

13 Sivaramakrishnan, S. et al. in The Oxford Handbook of the Auditory Brainstem (ed Karl Kandler) 0 (Oxford University Press, 2019).

14 Friauf, E. & Ostwald, J. Divergent projections of physiologically characterized rat ventral cochlear nucleus neurons as shown by intra-axonal injection of horseradish peroxidase. Exp Brain Res 73, 263–284 (1988). 10.1007/BF00248219

15 Hoffpauir, B. K., Grimes, J. L., Mathers, P. H. & Spirou, G. A. Synaptogenesis of the calyx of Held: rapid onset of function and one-to-one morphological innervation. J Neurosci 26, 5511–5523 (2006). 10.1523/JNEUROSCI.5525-05.2006

16 Holcomb, P. S. et al. Synaptic inputs compete during rapid formation of the calyx of Held: a new model system for neural development. J Neurosci 33, 12954–12969 (2013). 10.1523/JNEUROSCI.1087-13.2013

17 Kandler, K. & Friauf, E. Pre- and postnatal development of efferent connections of the cochlear nucleus in the rat. J Comp Neurol 328, 161–184 (1993). 10.1002/cne.903280202

18 Mikaelian, D. & Ruben, R. J. Development of Hearing in the Normal Cba-J Mouse:Correlation of Physiological Observations with Behavioral Responses and with Cochlear Anatomy. Acta Otolaryngolog 59, 451–461 (1965). 10.3109/00016486509124579

19 Erazo-Fischer, E., Striessnig, J. & Taschenberger, H. The role of physiological afferent nerve activity during in vivo maturation of the calyx of Held synapse. J Neurosci 27, 1725–1737 (2007). 10.1523/JNEUROSCI.4116-06.2007

20 Noh, J., Seal, R. P., Garver, J. A., Edwards, R. H. & Kandler, K. Glutamate co-release at GABA/glycinergic synapses is crucial for the refinement of an inhibitory map. Nat Neurosci 13, 232–238 (2010). 10.1038/nn.2478

21 Oleskevich, S., Youssoufian, M. & Walmsley, B. Presynaptic plasticity at two giant auditory synapses in normal and deaf mice. J Physiol 560, 709–719 (2004). 10.1113/jphysiol.2004.066662

22 Youssoufian, M., Oleskevich, S. & Walmsley, B. Development of a robust central auditory synapse in congenital deafness. J Neurophysiol 94, 3168–3180 (2005). 10.1152/jn.00342.2005

23 Babola, T. A. et al. Homeostatic Control of Spontaneous Activity in the Developing Auditory System. Neuron 99, 511–524 e515 (2018). 10.1016/j.neuron.2018.07.004

24 Kerschensteiner, D., Morgan, J. L., Parker, E. D., Lewis, R. M. & Wong, R. O. Neurotransmission selectively regulates synapse formation in parallel circuits in vivo. Nature 460, 1016–1020 (2009). 10.1038/nature08236

25 Kim, J. C. et al. Linking genetically defined neurons to behavior through a broadly applicable silencing allele. Neuron 63, 305–315 (2009). 10.1016/j.neuron.2009.07.010

26 Lessle, S. et al. Maintenance of a central high frequency synapse in the absence of synaptic activity. Front Cell Neurosci 18, 1404206 (2024). 10.3389/fncel.2024.1404206

27 Sando, R. et al. Assembly of Excitatory Synapses in the Absence of Glutamatergic Neurotransmission. Neuron 94, 312–321 e313 (2017). 10.1016/j.neuron.2017.03.047

28 Montesinos, M. S., Chen, Z. & Young, S. M., Jr. pUNISHER: a high-level expression cassette for use with recombinant viral vectors for rapid and long term in vivo neuronal expression in the CNS. J Neurophysiol 106, 3230–3244 (2011). 10.1152/jn.00713.2011

29 Schiavo, G. et al. Tetanus and botulinum-B neurotoxins block neurotransmitter release by proteolytic cleavage of synaptobrevin. Nature 359, 832–835 (1992). 10.1038/359832a0

30 Hoffpauir, B. K., Kolson, D. R., Mathers, P. H. & Spirou, G. A. Maturation of synaptic partners: functional phenotype and synaptic organization tuned in synchrony. J Physiol 588, 4365–4385 (2010). 10.1113/jphysiol.2010.198564

31 Hernandez, J. M. et al. Membrane fusion intermediates via directional and full assembly of the SNARE complex. Science 336, 1581–1584 (2012). 10.1126/science.1221976

32 Sierksma, M. C., Tedja, M. S. & Borst, J. G. In vivo matching of postsynaptic excitability with spontaneous synaptic inputs during formation of the rat calyx of Held synapse. J Physiol 595, 207–231 (2017). 10.1113/JP272780

33 Cant, N. B. & Gaston, K. C. Pathways connecting the right and left cochlear nuclei. J Comp Neurol 212, 313–326 (1982). 10.1002/cne.902120308

34 Needham, K. & Paolini, A. G. The commissural pathway and cochlear nucleus bushy neurons: an in vivo intracellular investigation. Brain Res 1134, 113–121 (2007). 10.1016/j.brainres.2006.11.058

35 Hsieh, C. Y. et al. Ephrin-B reverse signaling is required for formation of strictly contralateral auditory brainstem pathways. J Neurosci 30, 9840–9849 (2010). 10.1523/JNEUROSCI.0386-10.2010

36 Marrs, G. S. & Spirou, G. A. Embryonic assembly of auditory circuits: spiral ganglion and brainstem. J Physiol 590, 2391–2408 (2012). 10.1113/jphysiol.2011.226886

37 Borst, J. G. G., Helmchen, F. & Sakmann, B. Pre- and postsynaptic whole-cell recordings in the medial nucleus of the trapezoid body of the rat. Journal of Physiology 489.**3**, 825–840 (1995).

38 Forsythe, I. D. Direct patch recording from identified presynaptic terminals mediating glutamatergic EPSCs in the rat CNS, in vitro. J Physiol 479 **(Pt** **3**), 381–387 (1994). 10.1113/jphysiol.1994.sp020303

39 Rusu, S. I. & Borst, J. G. Developmental changes in intrinsic excitability of principal neurons in the rat medial nucleus of the trapezoid body. Dev Neurobiol 71, 284–295 (2011). 10.1002/dneu.20856

40 Geal-Dor, M., Freeman, S., Li, G. & Sohmer, H. Development of hearing in neonatal rats: air and bone conducted ABR thresholds. Hear Res 69, 236–242 (1993). 10.1016/0378-5955(93)90113-f

41 Meng, X. et al. Transient Subgranular Hyperconnectivity to L2/3 and Enhanced Pairwise Correlations During the Critical Period in the Mouse Auditory Cortex. Cereb Cortex 30, 1914–1930 (2020). 10.1093/cercor/bhz213

42 Tritsch, N. X., Yi, E., Gale, J. E., Glowatzki, E. & Bergles, D. E. The origin of spontaneous activity in the developing auditory system. Nature 450, 50–55 (2007). 10.1038/nature06233

43 Huberman, A. D., Feller, M. B. & Chapman, B. Mechanisms underlying development of visual maps and receptive fields. Annu Rev Neurosci 31, 479–509 (2008). 10.1146/annurev.neuro.31.060407.125533

44 Hua, Z. et al. v-SNARE composition distinguishes synaptic vesicle pools. Neuron 71, 474–487 (2011). 10.1016/j.neuron.2011.06.010

45 Ramirez, D. M., Khvotchev, M., Trauterman, B. & Kavalali, E. T. Vti1a identifies a vesicle pool that preferentially recycles at rest and maintains spontaneous neurotransmission. Neuron 73, 121–134 (2012). 10.1016/j.neuron.2011.10.034

46 Hamann, M., Billups, B. & Forsythe, I. D. Non-calyceal excitatory inputs mediate low fidelity synaptic transmission in rat auditory brainstem slices. Eur J Neurosci 18, 2899–2902 (2003). 10.1111/j.1460-9568.2003.03017.x

47 Banks, M. I., Pearce, R. A. & Smith, P. H. Hyperpolarization-activated cation current (Ih) in neurons of the medial nucleus of the trapezoid body: voltage-clamp analysis and enhancement by norepinephrine and cAMP suggest a modulatory mechanism in the auditory brain stem. J Neurophysiol 70, 1420–1432 (1993). 10.1152/jn.1993.70.4.1420

48 Koch, U., Braun, M., Kapfer, C. & Grothe, B. Distribution of HCN1 and HCN2 in rat auditory brainstem nuclei. Eur J Neurosci 20, 79–91 (2004). 10.1111/j.0953-816X.2004.03456.x

49 Banks, M. I. & Smith, P. H. Intracellular recordings from neurobiotin-labeled cells in brain slices of the rat medial nucleus of the trapezoid body. J Neurosci 12, 2819–2837 (1992). 10.1523/JNEUROSCI.12-07-02819.1992

50 Dodson, P. D., Barker, M. C. & Forsythe, I. D. Two heteromeric Kv1 potassium channels differentially regulate action potential firing. J Neurosci 22, 6953–6961 (2002). 20026709

51 Wang, L. Y., Gan, L., Forsythe, I. D. & Kaczmarek, L. K. Contribution of the Kv3.1 potassium channel to high-frequency firing in mouse auditory neurones. J Physiol 509 **(Pt** **1**), 183–194 (1998). 10.1111/j.1469-7793.1998.183bo.x

52 Hodgkin, A. L. & Huxley, A. F. Currents carried by sodium and potassium ions through the membrane of the giant axon of Loligo. J Physiol 116, 449–472 (1952). 10.1113/jphysiol.1952.sp004717

53 Leao, R. N., Svahn, K., Berntson, A. & Walmsley, B. Hyperpolarization-activated (I) currents in auditory brainstem neurons of normal and congenitally deaf mice. Eur J Neurosci 22, 147–157 (2005). 10.1111/j.1460-9568.2005.04185.x

54 Macica, C. M. et al. Modulation of the kv3.1b potassium channel isoform adjusts the fidelity of the firing pattern of auditory neurons. J Neurosci 23, 1133–1141 (2003). 10.1523/JNEUROSCI.23-04-01133.2003

55 Rothman, J. S. & Manis, P. B. The roles potassium currents play in regulating the electrical activity of ventral cochlear nucleus neurons. J Neurophysiol 89, 3097–3113 (2003). 10.1152/jn.00127.2002

56 Chequer Charan, D., et al. Volume electron microscopy reveals age-related circuit remodeling in the auditory brainstem. Front Cell Neurosci 16, 1070438 (2022). 10.3389/fncel.2022.1070438

57 Hsieh, C. Y. & Cramer, K. S. Deafferentation induces novel axonal projections in the auditory brainstem after hearing onset. J Comp Neurol 497, 589–599 (2006). 10.1002/cne.21002

58 Kitzes, L. M., Kageyama, G. H., Semple, M. N. & Kil, J. Development of ectopic projections from the ventral cochlear nucleus to the superior olivary complex induced by neonatal ablation of the contralateral cochlea. J Comp Neurol 353, 341–363 (1995). 10.1002/cne.903530303

59 Russell, F. A. & Moore, D. R. Afferent reorganisation within the superior olivary complex of the gerbil: development and induction by neonatal, unilateral cochlear removal. J Comp Neurol 352, 607–625 (1995). 10.1002/cne.903520409

60 Kuwabara, N., DiCaprio, R. A. & Zook, J. M. Afferents to the medial nucleus of the trapezoid body and their collateral projections. The Journal of Comparative Neurology 314, 684–706 (1991).

61 Spirou, G. A., Brownell, W. E. & Zidanic, M. Recordings from cat trapezoid body and HRP labeling of globular bushy cell axons. J Neurophysiol 63, 1169–1190 (1990). 10.1152/jn.1990.63.5.1169

62 Butts, D. A., Kanold, P. O. & Shatz, C. J. A burst-based “Hebbian” learning rule at retinogeniculate synapses links retinal waves to activity-dependent refinement. PLoS Biol 5, e61 (2007). 10.1371/journal.pbio.0050061

63 Nakashima, A. et al. Structured spike series specify gene expression patterns for olfactory circuit formation. Science 365 (2019). 10.1126/science.aaw5030

64 Kronander, E., Clark, C. & Schneggenburger, R. Role of BMP Signaling for the Formation of Auditory Brainstem Nuclei and Large Auditory Relay Synapses. Dev Neurobiol 79, 155–174 (2019). 10.1002/dneu.22661

65 Xiao, L. et al. BMP signaling specifies the development of a large and fast CNS synapse. Nat Neurosci 16, 856–864 (2013). 10.1038/nn.3414

66 Heller, D. T. et al. Astrocyte ensheathment of calyx-forming axons of the auditory brainstem precedes accelerated expression of myelin genes and myelination by oligodendrocytes. J Comp Neurol 532, e25552 (2024). 10.1002/cne.25552

67 Meng, X., Mukherjee, D., Kao, J. P. Y. & Kanold, P. O. Early peripheral activity alters nascent subplate circuits in the auditory cortex. Sci Adv 7 (2021). 10.1126/sciadv.abc9155

68 Muller, N. I. C., Sonntag, M., Maraslioglu, A., Hirtz, J. J. & Friauf, E. Topographic map refinement and synaptic strengthening of a sound localization circuit require spontaneous peripheral activity. J Physiol 597, 5469–5493 (2019). 10.1113/JP277757

69 Ebbers, L. et al. L-type Calcium Channel Cav1.2 Is Required for Maintenance of Auditory Brainstem Nuclei. J Biol Chem 290, 23692–23710 (2015). 10.1074/jbc.M115.672675

70 Hirtz, J. J. et al. Cav1.3 calcium channels are required for normal development of the auditory brainstem. J Neurosci 31, 8280–8294 (2011). 10.1523/JNEUROSCI.5098-10.2011

71 Satheesh, S. V. et al. Retrocochlear function of the peripheral deafness gene Cacna1d. Hum Mol Genet 21, 3896–3909 (2012). 10.1093/hmg/dds217

72 Brandebura, A. N. et al. Transcriptional profiling reveals roles of intercellular Fgf9 signaling in astrocyte maturation and synaptic refinement during brainstem development. J Biol Chem 298, 102176 (2022). 10.1016/j.jbc.2022.102176

73 Leao, R. N., Berntson, A., Forsythe, I. D. & Walmsley, B. Reduced low-voltage activated K+ conductances and enhanced central excitability in a congenitally deaf (dn/dn) mouse. J Physiol 559, 25–33 (2004). 10.1113/jphysiol.2004.067421

74 Clause, A. et al. The precise temporal pattern of prehearing spontaneous activity is necessary for tonotopic map refinement. Neuron 82, 822–835 (2014). 10.1016/j.neuron.2014.04.001

75 Di Guilmi, M. N. et al. Strengthening of the Efferent Olivocochlear System Leads to Synaptic Dysfunction and Tonotopy Disruption of a Central Auditory Nucleus. J Neurosci 39, 7037–7048 (2019). 10.1523/JNEUROSCI.2536-18.2019

76 Saul, S. M. et al. Math5 expression and function in the central auditory system. Mol Cell Neurosci 37, 153–169 (2008). 10.1016/j.mcn.2007.09.006

77 Jing, J. et al. Molecular logic for cellular specializations that initiate the auditory parallel processing pathways. Nat Commun 16, 489 (2025). 10.1038/s41467-024-55257-z

78 Altieri, S. C., Jalabi, W., Zhao, T., Romito-DiGiacomo, R. R. & Maricich, S. M. En1 directs superior olivary complex neuron positioning, survival, and expression of FoxP1. Dev Biol 408, 99–108 (2015). 10.1016/j.ydbio.2015.10.008

79 Brandebura, A. N. et al. Glial Cell Expansion Coincides with Neural Circuit Formation in the Developing Auditory Brainstem. Dev Neurobiol (2018). 10.1002/dneu.22633

80 Tong, H. et al. Regulation of Kv channel expression and neuronal excitability in rat medial nucleus of the trapezoid body maintained in organotypic culture. J Physiol 588, 1451–1468 (2010). 10.1113/jphysiol.2009.186676

81 Deisseroth, K., Bito, H. & Tsien, R. W. Signaling from synapse to nucleus: postsynaptic CREB phosphorylation during multiple forms of hippocampal synaptic plasticity. Neuron 16, 89–101 (1996). 10.1016/s0896-6273(00)80026-4

82 von Hehn, C. A., Bhattacharjee, A. & Kaczmarek, L. K. Loss of Kv3.1 tonotopicity and alterations in cAMP response element-binding protein signaling in central auditory neurons of hearing impaired mice. J Neurosci 24, 1936–1940 (2004). 10.1523/JNEUROSCI.4554-03.2004

83 Yap, E. L. & Greenberg, M. E. Activity-Regulated Transcription: Bridging the Gap between Neural Activity and Behavior. Neuron 100, 330–348 (2018). 10.1016/j.neuron.2018.10.013

84 Lujan, B., Dagostin, A. & von Gersdorff, H. Presynaptic Diversity Revealed by Ca(2+)-Permeable AMPA Receptors at the Calyx of Held Synapse. J Neurosci 39, 2981–2994 (2019). 10.1523/JNEUROSCI.2565-18.2019

85 Johnston, J., Griffin, S. J., Baker, C. & Forsythe, I. D. Kv4 (A-type) potassium currents in the mouse medial nucleus of the trapezoid body. Eur J Neurosci 27, 1391–1399 (2008). 10.1111/j.1460-9568.2008.06116.x

86 Mendonca, P. R. F., Kyle, V., Yeo, S. H., Colledge, W. H. & Robinson, H. P. C. Kv4.2 channel activity controls intrinsic firing dynamics of arcuate kisspeptin neurons. J Physiol 596, 885–899 (2018). 10.1113/JP274474

87 Johnston, J., Forsythe, I. D. & Kopp-Scheinpflug, C. Going native: voltage-gated potassium channels controlling neuronal excitability. J Physiol 588, 3187–3200 (2010). 10.1113/jphysiol.2010.191973

88 Montesinos, M. S., Satterfield, R. & Young, S. M., Jr. Helper-Dependent Adenoviral Vectors and Their Use for Neuroscience Applications. Methods Mol Biol 1474, 73–90 (2016). 10.1007/978-1-4939-6352-2_5

89 Keine, C., Al-Yaari, M., Radulovic, T. & Young, S. M., Jr. Stereotactic Delivery of Helper-dependent Adenoviral Viral Vectors at Distinct Developmental Time Points to Perform Age-dependent Molecular Manipulations of the Mouse Calyx of Held. Bio Protoc 13, e4793 (2023). 10.21769/BioProtoc.4793

90 Leao, R. N. et al. Topographic organization in the auditory brainstem of juvenile mice is disrupted in congenital deafness. J Physiol 571, 563–578 (2006). 10.1113/jphysiol.2005.098780

91 Awatramani, G. B., Turecek, R. & Trussell, L. O. Staggered development of GABAergic and glycinergic transmission in the MNTB. J Neurophysiol 93, 819–828 (2005). 10.1152/jn.00798.2004

92 Kushmerick, C., Renden, R. & von Gersdorff, H. Physiological temperatures reduce the rate of vesicle pool depletion and short-term depression via an acceleration of vesicle recruitment. J Neurosci 26, 1366–1377 (2006). 10.1523/JNEUROSCI.3889-05.2006

93 Clements, J. D. & Bekkers, J. M. Detection of spontaneous synaptic events with an optimally scaled template. Biophys J 73, 220–229 (1997). 10.1016/S0006-3495(97)78062-7

94 Jonas, P., Major, G. & Sakmann, B. Quantal components of unitary EPSCs at the mossy fibre synapse on CA3 pyramidal cells of rat hippocampus. J Physiol 472, 615–663 (1993). 10.1113/jphysiol.1993.sp019965

95 Hines, M. L. & Carnevale, N. T. NEURON: a tool for neuroscientists. Neuroscientist 7, 123–135 (2001). 10.1177/107385840100700207

