## Supplemental Figures and Tables for "Pre-Sensory Spontaneous Activity Accelerates Coordinated Maturation of Synaptic Partners and Drives Transition to the Mature Physiological Phenotype"

**This document includes:**

Figures S1 to S4
Tables S1 to S12
SI References

### Supplementary Figures and Tables

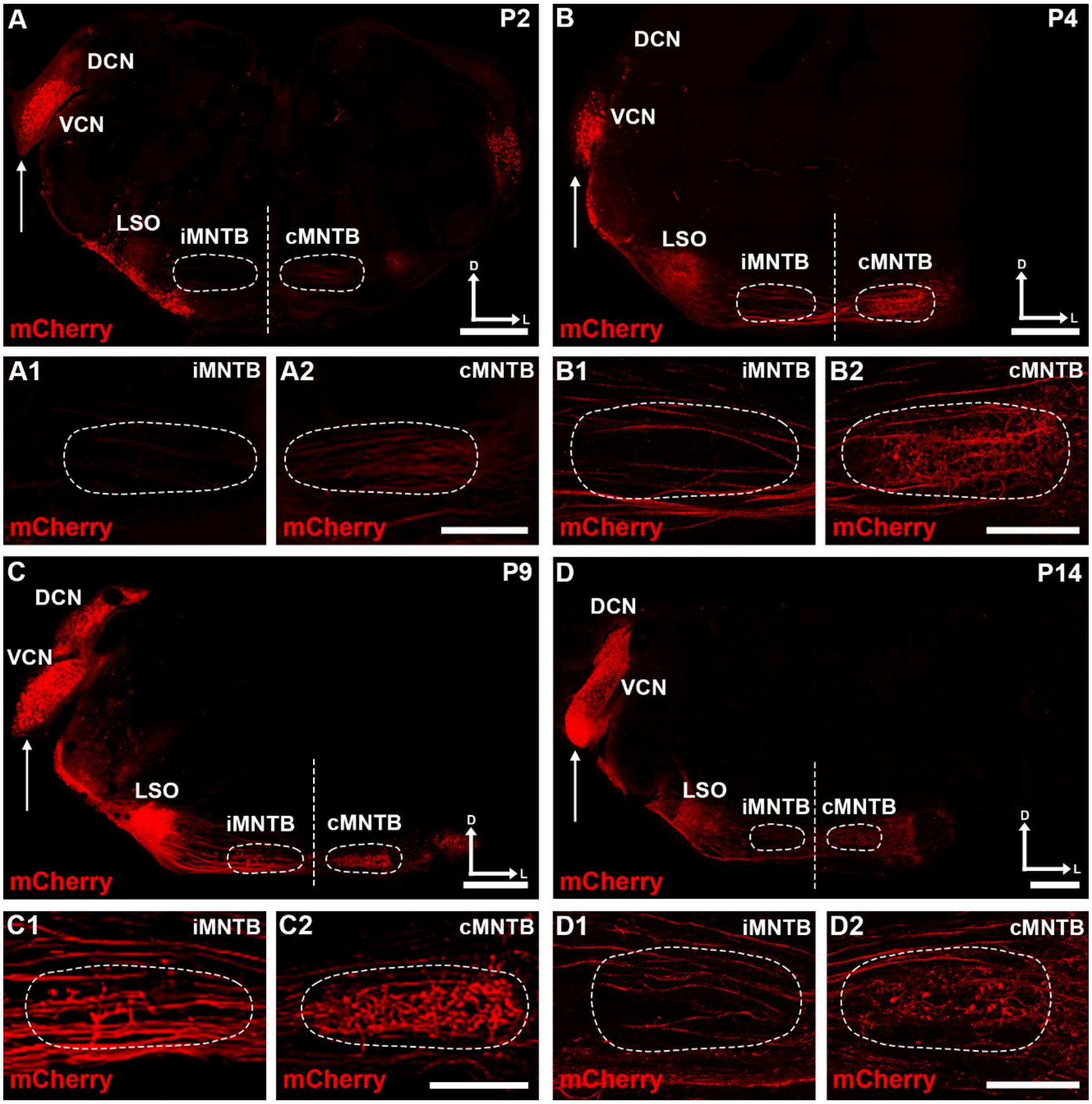


**Figure S1.** Expression pattern following unilateral viral vector injection into the ventral cochlear nucleus (VCN) across development. (A-D), Injection of the dual expression HdAd vector (mCherry reporter molecule co-expressed with TeNT light chain) into the VCN was performed at P0 followed by transcardial perfusion with 4% PFA at P2, P4, P9, and P14. Endogenous mCherry labeling was amplified with primary antibodies against dsRed following standard immunohistochemistry procedures. Arrows indicate the viral injection site. Vertical dashed line indicates the midline. The ipsilateral and contralateral MNTB (iMNTB and cMNTB, respectively), to the injection site is outlined in a dashed line. Labeling in the injected cochlear nucleus is confined to the neuronal cell bodies in VCN, with afferent projection fibers to the dorsal cochlear nucleus (DCN) and lateral superior olive (LSO) labeled. Dorsal and lateral axes are indicated above scale bar. Scale bar: 500 μm. (A1-D1 and A2-D2), High-magnification images of iMNTB and cMNTB. Scale bar: 200 μm.

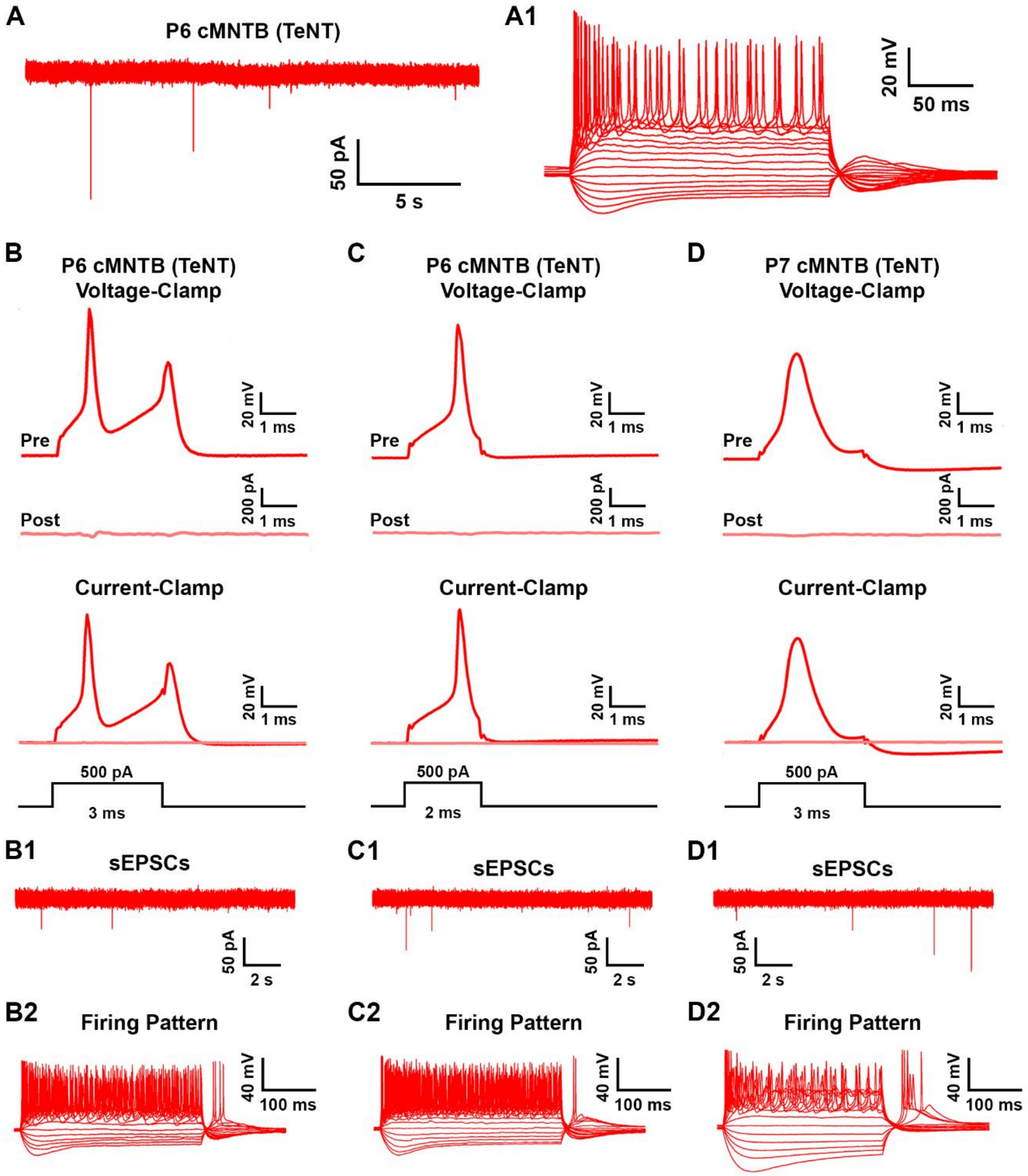


**Figure S2.** Simultaneous paired electrophysiological recordings from the cMNTB following TeNT expression. (A), Exemplary voltage-clamp trace of the P6 cMNTB PN from Figure 1E3 showing sEPSCs consistent with P6 TeNT electrophysiological data. The cell was clamped at a holding potential of -73 mV. Recordings were made after bath application of the inhibitory synaptic blockers Gabazine (GABAA receptor antagonist, 10 μM) and strychnine (glycine receptor antagonist, 2 μM). (A1), Exemplary current-clamp trace of the P6 cMNTB PN from Figure 1E3 showing a tonic firing pattern consistent with P6 TeNT electrophysiological data. Data was recorded at resting membrane potential and shown as a −100 pA step with successive steps at 20 pA increments. (B-D), Additional paired recordings, at P6 and P7, from the cMNTB showing abolished evoked neurotransmission. Evoked responses in PNs, following a 2-3 ms, 500 pA depolarizing current injection via the presynaptic recording pipette, in voltage (EPSC, top) and current-clamp (action potential, bottom) configuration. (B1-D1 and B2-D2), Exemplary voltage and current-clamp traces from the cMNTB PNs corresponding to the paired recording in panels (B-D), respectively, showing reduced frequency of sEPSCs and characteristic tonic firing pattern phenotype.

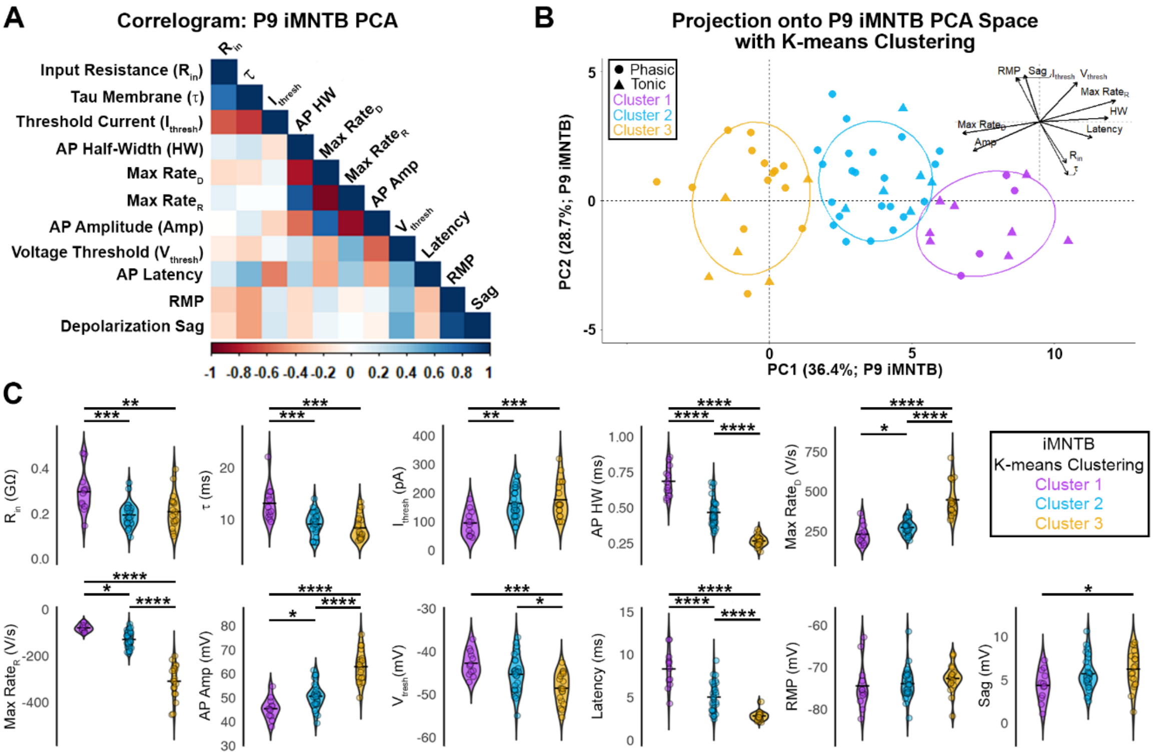


**Figure S3.** Principal component analysis (PCA) of electrophysiological parameters comparing the maturational trajectory of iMNTB PNs (without TeNT expression). (A), Correlogram showing the correlation between the electrophysiological parameters and the first two principal components for P9 iMNTB PNs. The color and intensity represent the degree of correlation, with the scale shown at the bottom of the plot. (B), K-means algorithm, with three clusters shown by different colors, was applied to PCA projection in Figure 6A to identify cells with similar physiological profiles and shows the maturational profile in PCA space from lower right (purple) to upper left (gold). Inset shows the loading vectors from Figure 6A. (C), Violin plots of the 11 electrophysiological parameters in each cluster shown for panel (B), demonstrating the maturational progression of parameter values in iMNTB PCA space. Each color corresponds to the clusters identified from the K-means algorithm. Significance levels are indicated according to the convention: *P < 0.05, **P < 0.01, ***P < 0.001, ****P < 0.0001, and not significant (ns). Statistical results for panel (C) are reported in Table S10.

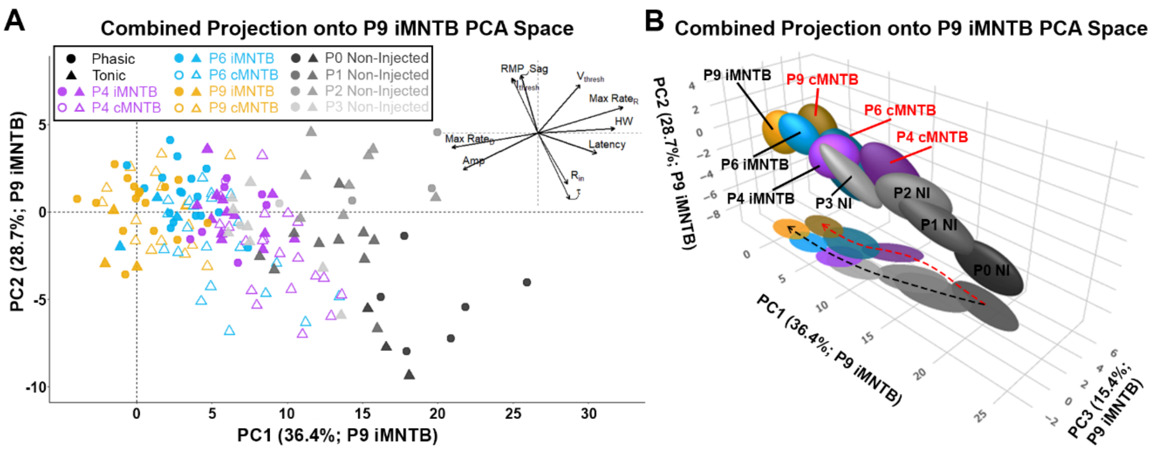


**Figure S4.** Principal component analysis (PCA) of electrophysiological parameters comparing the physiological profiles of PNs with (cMNTB) and without (iMNTB) TeNT expression to non-injected control PNs (A), Combined projection of all groups (P4 and P6 iMNTB and P4, P6, and P9 cMNTB PNs) and electrophysiological recordings from non-injected mice (P0 (n = 9 cells), P1 (n = 10 cells), P2 (n = 10 cells), and P3 (n = 10 cells)) onto the 2-dimensional PCA space of P9 iMNTB. Circles and triangles represent phasic and tonic firing patterns, respectively. Filled symbols correspond to iMNTB and non-injected control PNs and open symbols correspond to transduced cMNTB PNs (TeNT expression). Inset shows the loading vectors from PCA of P9 iMNTB PNs presented in Figure 6A. (B), 3D plot showing ellipsoids that are centered and scaled (one standard deviation) for each experimental group (P4, P6, and P9 iMNTB and cMNTB PNs) and non-injected PNs (P0, P1, P2, and P3). Transduced (cMNTB) PNs follow a different maturation trajectory than non-transduced (iMNTB) PNs with a divergence from early postnatal non-injected PNs. Different maturation trajectories (black and red dashed arrows for iMNTB and cMNTB, respectively) are also visualized by projection of ellipsoids onto the PC1 and PC3 2D plane.

**Table S1.** Quantification of mCherry labeling (co-expressed with TeNT) in the MNTB

| Age/Experimental Condition | Large Terminal mCherry+/Vglut+ | Large Terminal mCherry-/Vglut+ | Punctate Labeling (mCherry+) | Punctate Labeling (mCherry-) |
|---|---|---|---|---|
| P4 iMNTB (Control; n = 2 brains) | 0 (0%) | 219 (90.5%) | 0 (0%) | 23 (9.5%) |
| P4 cMNTB (TeNT expression; n = 2 brains) | 40 (12.4%) | 30 (9.3 %) | 251 (77.7%) | 2 (0.6%) |
| P6 iMNTB (Control; n = 3 brains) | 1 (0.3%) | 303 (93.5%) | 0 (0%) | 20 (6.2%) |
| P6 cMNTB (TeNT expression; n = 3 brains) | 234 (77.2%) | 13 (4.3%) | 52 (17.2%) | 4 (1.3%) |
| P9 iMNTB (Control; n = 2 brains) | 18 (6.6%) | 248 (91.2%) | 0 (0%) | 6 (2.2%) |
| P9 cMNTB (TeNT expression; n = 2 brains) | 127 (83.6%) | 12 (7.9%) | 13 (8.6%) | 0 (0%) |
| P14 iMNTB (Control; n = 2 brains) | 2 (0.9%) | 230 (98.7%) | 0 (0%) | 1 (0.4%) |
| P14 cMNTB (TeNT expression; n = 2 brains) | 109 (63.7%) | 50 (29.2%) | 9 (5.3%) | 3 (1.8%) |

*Data presented as absolute numbers with the total percentage in parentheses.*

**Table S2.** Parameters for MNTB PN sEPSCs following TeNT expression

| MNTB PN Age | Amplitude (pA) | Decay Time Constant, $\tau$ (ms) | Risetime 20-80% (ms) | Frequency (Hz) |
|---|---|---|---|---|
| P4 Non-injected/Control (n = 10) | 70.41 ± 11.81 | 0.90 ± 0.12 | 0.14 ± 0.01 | 2.75 ± 1.63 |
| P4 iMNTB/Control (n = 20) | 65.94 ± 11.45 | 0.76 ± 0.14 | 0.14 ± 0.02 | 2.14 ± 1.76 |
| P4 cMNTB/TeNT (n = 21) | 55.04 ± 12.12 | 1.14 ± 0.34 | 0.25 ± 0.10 | 0.53 ± 0.66 |
| P6 Non-injected/Control (n = 15) | 68.51 ± 9.87 | 0.66 ± 0.10 | 0.13 ± 0.003 | 3.07 ± 3.45 |
| P6 iMNTB/Control (n = 20) | 62.68 ± 14.28 | 0.72 ± 0.18 | 0.13 ± 0.01 | 2.50 ± 1.94 |
| P6 cMNTB/TeNT (n = 21) | 61.54 ± 15.13 | 1.10 ± 0.27 | 0.20 ± 0.06 | 0.60 ± 0.40 |
| P9 Non-injected/Control (n = 13) | 48.63 ± 11.81 | 0.47 ± 0.12 | 0.13 ± 0.006 | 2.52 ± 1.61 |
| P9 iMNTB/Control (n = 19) | 48.00 ± 8.86 | 0.47 ± 0.16 | 0.13 ± 0.03 | 0.97 ± 1.35 |
| P9 cMNTB/TeNT (n = 17) | 63.54 ± 18.44 | 1.04 ± 0.31 | 0.16 ± 0.03 | 0.83 ± 0.50 |
| P14 Non-injected/Control (n = 12) | 61.21 ± 10.80 | 0.24 ± 0.08 | 0.11 ± 0.005 | 5.65 ± 4.50 |
| P14 iMNTB/Control (n = 15) | 67.86 ± 22.24 | 0.29 ± 0.18 | 0.12 ± 0.01 | 4.60 ± 6.41 |
| P14 cMNTB/TeNT (n = 11) | 68.30 ± 18.80 | 0.60 ± 0.14 | 0.13 ± 0.01 | 1.33 ± 1.05 |

*Data presented as means ± standard deviation based on the average values for individual cells.*
*n: number of cells.*

**Table S3.** Parameters for MNTB PN biophysical properties following TeNT expression

| MNTB PN Age | Slope Resistance (GΩ) | $\tau_{Membrane}$ (ms) | RMP (mV) | Sag (mV) |
|---|---|---|---|---|
| P4 iMNTB (n = 20) | 0.25 ± 0.09 | 11.35 ± 3.29 | -74.32 ± 5.61 | 5.00 ± 2.21 |
| P4 cMNTB (n = 21) | 0.45 ± 0.13 | 18.95 ± 5.92 | -68.54 ± 4.77 | 6.28 ± 3.12 |
| P6 iMNTB (n = 20) | 0.2 ± 0.07 | 9.01 ± 2.83 | -73.26 ± 2.43 | 6.04 ± 1.70 |
| P6 cMNTB (n = 21) | 0.34 ± 0.15 | 15.29 ± 6.5 | -71.50 ± 4.45 | 5.20 ± 2.72 |
| P9 iMNTB (n = 19) | 0.21 ± 0.7 | 8.69 ± 2.49 | -73.24 ± 3.51 | 5.88 ± 2.24 |
| P9 cMNTB (n = 17) | 0.23 ± 0.09 | 11.73 ± 4.02 | -68.77 ± 4.08 | 6.81 ± 2.38 |
| P14 iMNTB (n = 15) | 0.22 ± 0.10 | 8.41 ± 2.64 | -72.53 ± 2.67 | 5.39 ± 1.64 |
| P14 cMNTB (n = 11) | 0.30 ± 0.11 | 11.31 ± 3.15 | -70.44 ± 5.92 | 5.66 ± 3.20 |

*Data presented as means ± standard deviation based on the average values for individual cells.*
*n: number of cells.*

**Table S4.** Parameters for MNTB PN AP waveform kinetics following TeNT expression

| MNTB PN Age | Latency (ms) | $I_{Threshold}$ (pA) | Amplitude (mV) | Half-Width (ms) | $V_{Threshold}$ (mV) | $Rate_{Depol}$ (V/s) | $Rate_{Repol}$ (V/s) |
|---|---|---|---|---|---|---|---|
| P4 iMNTB (n = 20) | 7.41 ± 2.01 | 120.75 ± 42.56 | 48.03 ± 5.42 | 0.63 ± 0.12 | -43.17 ± 3.46 | 263.68 ± 68.99 | -84.71 ± 20.21 |
| P4 cMNTB (n = 21) | 10.92 ± 5.77 | 46.43 ± 37.65 | 50.03 ± 5.91 | 0.75 ± 0.15 | -41.49 ± 4.83 | 201.53 ± 53.98 | -74.96 ± 16.28 |
| P6 iMNTB (n = 20) | 4.34 ± 1.75 | 172.75 ± 65.24 | 53.18 ± 7.00 | 0.42 ± 0.09 | -45.94 ± 3.96 | 295.84 ± 89.52 | -160.88 ± 64.63 |
| P6 cMNTB (n = 21) | 8.26 ± 2.24 | 68.10 ± 29.60 | 57.26 ± 7.74 | 0.54 ± 0.24 | -43.83 ± 4.75 | 312.68 ± 101.97 | -126.82 ± 39.47 |
| P9 iMNTB (n = 19) | 3.18 ± 1.57 | 170.26 ± 64.30 | 59.86 ± 9.50 | 0.28 ± 0.06 | -48.43 ± 3.94 | 410.74 ± 129.89 | -288.85 ± 90.85 |
| P9 cMNTB (n = 17) | 5.67 ± 1.61 | 104.12 ± 43.88 | 57.56 ± 10.87 | 0.32 ± 0.07 | -42.86 ± 4.21 | 401.14 ± 143.39 | -227.19 ± 76.47 |
| P14 iMNTB (n = 15) | 2.81 ± 0.47 | 121.33 ± 56.55 | 60.69 ± 5.34 | 0.22 ± 0.04 | -52.87 ± 2.32 | 452.75 ± 115.38 | -400.77± 117.39 |
| P14 cMNTB (n = 11) | 3.63 ± 1.06 | 70.00 ± 52.35 | 70.19 ± 7.85 | 0.19 ± 0.04 | -51.73 ± 3.99 | 640.99 ± 139.83 | -518.62 ± 144.54 |

*Data presented as means ± standard deviation based on the average values for individual cells.*
*n: number of cells.*

**Table S5.** Parameters for CH and MNTB PN morphology following TeNT expression

| MNTB PN Age | CH Volume (μm) | CH Thickness (μm) | PN Volume ($\mu m^3$) | PN Surface Area ($\mu m^2$) |
|---|---|---|---|---|
| P4 iMNTB (n = 83) | 572.06 ± 274.21 | 1.45 ± 0.47 | 2077.22 ± 419.01 | 783.81 ± 105.32 |
| P4 cMNTB (n = 33) | 481.13 ± 252.60 | 2.43 ± 0.82 | 1849.11 ± 345.39 | 725.71 ± 91.80 |
| P6 iMNTB (n = 107) | 1070.59 ± 505.23 | 1.29 ± 0.44 | 2101.88 ± 423.59 | 790.09 ± 105.03 |
| P6 cMNTB (n = 87) | 737.37 ± 449.97 | 2.30 ± 0.95 | 1975.74 ± 346.72 | 758.94 ± 88.05 |
| P9 iMNTB (n = 96) | 1185.82 ± 474.88 | 1.19 ± 0.31 | 2141.48 ± 368.40 | 802.63 ± 91.43 |
| P9 cMNTB (n = 55) | 551.93 ± 331.97 | 2.24 ± 1.16 | 2112.32 ± 323.05 | 790.70 ± 80.94 |
| P14 iMNTB (n = 98) | 1401.31 ± 620.42 | 0.91 ± 0.23 | 2461.07 ± 655.96 | 874.90 ± 154.03 |
| P14 cMNTB (n = 40) | 649.24 ± 428.55 | 2.30 ± 1.33 | 2359.73 ± 472.82 | 853.58 ± 112.61 |

*Data presented as means ± standard deviation based on the average values for individual cells. n: number of cells.*

**Table S6.** Statistics corresponding to results in Figure 2B-E.

| | MNTB PN Comparison | Amplitude | Decay Time Constant | Risetime | Frequency |
|---|---|---|---|---|---|
| **P4** | Group Comparison | One-way ANOVA: P = 1.7E-3 | One-way ANOVA: P < 0.0001 | Kruskal-Wallis: P < 0.0001 | Kruskal-Wallis: P < 0.0001 |
| | Control vs. iMNTB | P = 0.60 | P = 0.30 | P > 0.99 | P > 0.99 |
| | Control vs. cMNTB | P = 3.9E-3 | P = 3.6E-2 | P = 1.4E-2 | P < 0.0001 |
| | iMNTB vs. cMNTB | P = 1.3E-2 | P < 0.0001 | P = 2.0E-4 | P = 0.0001 |
| **P6** | Group Comparison | Kruskal-Wallis: P = 7.2E-2 | One-way ANOVA: P < 0.0001 | Kruskal-Wallis: P < 0.0001 | Kruskal-Wallis: P < 0.0001 |
| | Control vs. iMNTB | P = 0.20 | P = 0.60 | P > 0.99 | P > 0.99 |
| | Control vs. cMNTB | P = 0.09 | P < 0.0001 | P = 2.0E-4 | P < 0.0001 |
| | iMNTB vs. cMNTB | P > 0.99 | P < 0.0001 | P < 0.0001 | P < 0.0001 |
| **P9** | Group Comparison | One-way ANOVA: P = 9.0E-4 | One-way ANOVA: P < 0.0001 | Kruskal-Wallis: P < 0.0001 | Kruskal-Wallis: P = 8.0E-4 |
| | Control vs. iMNTB | P = 0.99 | P = 0.99 | P > 0.99 | P = 6.0E-4 |
| | Control vs. cMNTB | P = 6.4E-3 | P < 0.0001 | P = 2.0E-4 | P = 1.7E-2 |
| | iMNTB vs. cMNTB | P = 1.6E-3 | P < 0.0001 | P = 3.0E-4 | P > 0.99 |
| **P14** | Group Comparison | One-way ANOVA: P = 0.83 | One-way ANOVA: P < 0.0001 | One-way ANOVA: P = 2.0E-4 | Kruskal-Wallis: P = 4.1E-2 |
| | Control vs. iMNTB | P = 0.83 | P = 0.31 | P = 0.22 | P = 0.27 |
| | Control vs. cMNTB | P = 0.90 | P < 0.0001 | P = 2.0E-4 | P = 3.9E-2 |
| | iMNTB vs. cMNTB | P = 0.99 | P < 0.0001 | P = 8.3E-3 | P > 0.99 |

*Tukey's and Dunn's test for multiple comparisons was utilized following one-way ANOVA and Kruskal-ANOVA, respectively (based on the results of a Shapiro-Wilk normality test).*

**Table S7.** Statistics corresponding to results in Figure 3B-E.

| **MNTB PN Comparison** | **Slope Resistance** | $\tau_{Membrane}$ | **RMP** | **Sag** |
|---|---|---|---|---|
| P4 Control vs. TeNT | Mann-Whitney: P < 0.0001 | Mann-Whitney: P < 0.0001 | Mann-Whitney: P = 4.0E-4 | Unpaired t-test: P = 0.14 |
| P6 Control vs. TeNT | Unpaired t-test: P = 6.0E-4 | Mann-Whitney: P = 0.0001 | Unpaired t-test: P = 0.13 | Unpaired t-test: P = 0.25 |
| P9 Control vs. TeNT | Unpaired t-test: P = 0.38 | Unpaired t-test: P = 9.3E-3 | Unpaired t-test: P = 1.2E-3 | Unpaired t-test: P = 0.24 |
| P14 Control vs. TeNT | Mann-Whitney: P = 4.1E-2 | Mann-Whitney: P 0.02 | Unpaired t-test: P = 0.24 | Unpaired t-test: P = 0.78 |

*An unpaired t-test or Mann-Whitney Test was run following the results of a Shapiro-Wilk normality test.*

**Table S8.** Statistics corresponding to results in Figure 4C-I.

| MNTB PN Comparison | Latency | $I_{Threshold}$ | Amplitude | Half-Width | $V_{Threshold}$ | $Rate_{Depol}$ | $Rate_{Repol}$ |
|---|---|---|---|---|---|---|---|
| P4 Control vs. TeNT | Mann-Whitney: P = 1.2E-2 | Mann-Whitney: P < 0.0001 | Unpaired t-test: P = 0.27 | Unpaired t-test: P = 6.8E-3 | Unpaired t-test: P = 0.21 | Unpaired t-test: P = 2.6E-3 | Unpaired t-test: P = 9.6E-2 |
| P6 Control vs. TeNT | Mann-Whitney: P < 0.0001 | Unpaired t-test: P < 0.0001 | Unpaired t-test: P = 8.5E-2 | Mann-Whitney: P = 8.3E-2 | Unpaired t-test: P = 0.13 | Mann-Whitney: P = 0.61 | Mann-Whitney: P = 0.10 |
| P9 Control vs. TeNT | Mann-Whitney: P < 0.0001 | Unpaired t-test: P = 1.1E-3 | Unpaired t-test: P = 0.50 | Unpaired t-test: P = 8.9E-2 | Unpaired t-test: P = 2.0E-4 | Unpaired t-test: P = 0.83 | Unpaired t-test: P = 3.6E-2 |
| P14 Control vs. TeNT | Unpaired t-test: P = 1.3E-2 | Unpaired t-test: P = 2.7E-2 | Unpaired t-test: P = 1.2E-3 | Unpaired t-test: P = 6.6E-2 | Unpaired t-test: P = 0.37 | Unpaired t-test: P = 1.0E-3 | Unpaired t-test: P = 3.1E-2 |

*An unpaired t-test or Mann-Whitney Test was run following the results of a Shapiro-Wilk normality test.*

**Table S9.** Statistics corresponding to results in Figure 7K-M.

| MNTB PN Comparison | CH Volume | CH Thickness | PN Volume | PN Surface Area |
|---|---|---|---|---|
| P4 Control vs. TeNT | Mann-Whitney: P = 0.12 | Mann-Whitney: P < 0.0001 | Unpaired t-test: P = 6.5E-3 | Unpaired t-test: P = 6.4E-3 |
| P6 Control vs. TeNT | Mann-Whitney: P < 0.0001 | Mann-Whitney: P < 0.0001 | Mann-Whitney: P = 3.4E-2 | Mann-Whitney: P = 3.4E-2 |
| P9 Control vs. TeNT | Mann-Whitney: P < 0.0001 | Mann-Whitney: P < 0.0001 | Unpaired t-test: P = 0.63 | Unpaired t-test: P = 0.43 |
| P14 Control vs. TeNT | Mann-Whitney: P < 0.0001 | Mann-Whitney: P < 0.0001 | Mann-Whitney: P = 0.76 | Mann-Whitney: P = 0.76 |

*An unpaired t-test or Mann-Whitney Test was run following the results of a Shapiro-Wilk normality test.*

**Table S10.** Statistics corresponding to results in Figure S3C for iMNTB PNs.

| Parameter | Group Comparison | Cluster 1 vs. 2 | Cluster 1 vs. 3 | Cluster 2 vs. 3 |
|---|---|---|---|---|
| Input Resistance | One-way ANOVA: P = 3.8E-4 | Unpaired t-test: P = 2.9E-4 | Unpaired t-test: P = 3.8E-3 | Unpaired t-test: P = 0.78 |
| $\tau_{Membrane}$ | Kruskal-Wallis: P = 2.9E-4 | Wilcoxon Test: P = 7.0E-4 | Wilcoxon Test: P = 2.9E-4 | Wilcoxon Test: P = 0.32 |
| Threshold Current | One-way ANOVA: P = 4.9E-4 | Unpaired t-test: P = 2.0E-3 | Unpaired t-test: P = 5.8E-4 | Unpaired t-test: P = 0.72 |
| AP Half-Width | One-way ANOVA: P < 0.0001 | Unpaired t-test: P < 0.0001 | Unpaired t-test: P < 0.0001 | Unpaired t-test: P < 0.0001 |
| Max Rate$_D$ | Kruskal-Wallis: P < 0.0001 | Wilcoxon Test: P = 2.4E-2 | Wilcoxon Test: P < 0.0001 | Wilcoxon Test: P < 0.0001 |
| Max Rate$_R$ | One-way ANOVA: P < 0.0001 | Unpaired t-test: P = 1.2E-2 | Unpaired t-test: P < 0.0001 | Unpaired t-test: P < 0.0001 |
| AP Amplitude | One-way ANOVA: P < 0.0001 | Unpaired t-test: P = 2.6E-2 | Unpaired t-test: P < 0.0001 | Unpaired t-test: P < 0.0001 |
| Voltage Threshold | One-way ANOVA: P = 2.8E-4 | Unpaired t-test: P = 0.13 | Unpaired t-test: P = 3.4E-4 | Unpaired t-test: P = 1.6E-2 |
| Latency | Kruskal-Wallis: P < 0.0001 | Wilcoxon Test: P < 0.0001 | Wilcoxon Test: P < 0.0001 | Wilcoxon Test: P < 0.0001 |
| RMP | Kruskal-Wallis: P = 0.09 | N/A | N/A | N/A |
| Sag | One-way ANOVA: P = 4.2E-2 | Unpaired t-test: P = 0.13 | Unpaired t-test: P = 3.5E-2 | Unpaired t-test: P = 0.64 |

*An unpaired t-test or Wilcoxon test for multiple comparisons was utilized following one-way ANOVA and Kruskal-ANOVA, respectively (based on the results of a Shapiro-Wilk normality test).*

**Table S11.** PERMANOVA results for each experimental group corresponding to Figure S4.

| Group 1 | Group 2 | F-value | P-value |
|---|---|---|---|
| | P1 NI | 6.56 | 4.50E-3 |
| | P2 NI | 14.33 | 3.00E-4 |
| | P4 TeNT | 40.57 | 1.00E-4 |
| | P3 NI | 42.40 | 1.00E-4 |
| | P6 TeNT | 63.28 | 1.00E-4 |
| P0 NI | P4 iMNTB | 104.37 | 1.00E-4 |
| | P9 TeNT | 138.26 | 1.00E-4 |
| | P6 iMNTB | 170.73 | 1.00E-4 |
| | P9 iMNTB | 198.60 | 1.00E-4 |
| | P2 NI | 1.83 | 0.16 |
| | P3 NI | 17.21 | 5.00E-4 |
| | P4 TeNT | 16.69 | 1.00E-4 |
| P1 NI | P6 TeNT | 34.83 | 1.00E-4 |
| | P4 iMNTB | 51.18 | 1.00E-4 |
| | P9 TeNT | 88.70 | 1.00E-4 |
| | P6 iMNTB | 103.10 | 1.00E-4 |
| | P9 iMNTB | 133.43 | 1.00E-4 |
| | P3 NI | 8.63 | 2.40E-3 |
| | P4 TeNT | 6.73 | 2.10E-3 |
| | P6 TeNT | 19.56 | 1.00E-4 |
| P2 NI | P4 iMNTB | 29.56 | 1.00E-4 |
| | P9 TeNT | 61.40 | 1.00E-4 |
| | P6 iMNTB | 71.37 | 1.00E-4 |
| | P9 iMNTB | 98.63 | 1.00E-4 |
| | P4 iMNTB | 2.97 | 0.04 |
| | P6 TeNT | 5.08 | 7.70E-3 |
| P3 NI | P4 TeNT | 6.64 | 3.30E-3 |
| | P6 iMNTB | 22.80 | 1.00E-4 |
| | P9 TeNT | 25.57 | 1.00E-4 |
| | P9 iMNTB | 42.92 | 1.00E-4 |
| | P6 TeNT | 5.74 | 9.00E-4 |
| | P6 iMNTB | 16.48 | 1.00E-4 |
| P4 iMNTB | P9 TeNT | 24.19 | 1.00E-4 |
| | P9 iMNTB | 44.12 | 1.00E-4 |
| | P6 TeNT | 9.05 | 8.00E-4 |
| | P4 iMNTB | 19.82 | 1.00E-4 |
| | P9 TeNT | 41.38 | 1.00E-4 |
| P4 TeNT | P6 iMNTB | 54.32 | 1.00E-4 |
| | P9 iMNTB | 79.37 | 1.00E-4 |
| | P9 TeNT | 7.91 | 4.00E-4 |
| P6 iMNTB | P9 iMNTB | 9.30 | 1.00E-4 |
| | P9 TeNT | 10.34 | 2.00E-4 |
| | P6 iMNTB | 14.85 | 1.00E-4 |
| P6 TeNT | P9 iMNTB | 26.10 | 1.00E-4 |
| | P9 iMNTB | 7.50 | 1.00E-4 |
| P9 TeNT | P9 iMNTB | 7.50 | 1.00E-4 |

*Abbreviations: NI, Non-injected.*

**Table S12.** Parameters for the kinetics of $Na_V$, $KHT_V$, $KLT_V$, and HCN

| | ***$Na_V$* Activation** | ***$Na_V$* Inactivation** | ***$KHT_V$* Activation** | ***$KHT_V$* Inactivation** | ***$KLT_V$* Activation** | ***HCN* Activation** |
|---|---|---|---|---|---|---|
| | ***m*** | ***h*** | ***n*** | ***j*** | ***o*** | ***u*** |
| $C_\alpha$ ($ms^{-1}$) | 76.6 | 0.00013 | 0.2719 | 0.00713 | 1.2 | $9.12 \times 10^{-8}$ |
| $k_\alpha$ ($mV^{-1}$) | 0.37 | -0.1216 | 0.04 | -0.1942 | 0.03512 | -0.01 |
| $C_\beta$ ($ms^{-1}$) | 6.90852 | 1.999 | 0.1974 | 0.0935 | 0.2248 | $2.1 \times 10^{-3}$ |
| $k_\beta$ ($mV^{-1}$) | -0.043 | 0.0384 | 0 | 0.0058 | -0.0319 | 0 |

*Parameters for voltage dependence and kinetics were based on previous models of PNs[1-5].*

### SI References


1 Leao, R. N., Svahn, K., Berntson, A. & Walmsley, B. Hyperpolarization-activated (I) currents in auditory brainstem neurons of normal and congenitally deaf mice. *Eur J Neurosci* **22**, 147-157 (2005). https://doi.org:10.1111/j.1460-9568.2005.04185.x

2 Macica, C. M. *et al.* Modulation of the kv3.1b potassium channel isoform adjusts the fidelity of the firing pattern of auditory neurons. *J Neurosci* **23**, 1133-1141 (2003). https://doi.org:10.1523/JNEUROSCI.23-04-01133.2003

3 Rothman, J. S. & Manis, P. B. The roles potassium currents play in regulating the electrical activity of ventral cochlear nucleus neurons. *J Neurophysiol* **89**, 3097-3113 (2003). https://doi.org:10.1152/jn.00127.2002

4 Sierksma, M. C., Tedja, M. S. & Borst, J. G. In vivo matching of postsynaptic excitability with spontaneous synaptic inputs during formation of the rat calyx of Held synapse. *J Physiol* **595**, 207-231 (2017). https://doi.org:10.1113/JP272780

5 Wang, L. Y., Gan, L., Forsythe, I. D. & Kaczmarek, L. K. Contribution of the Kv3.1 potassium channel to high-frequency firing in mouse auditory neurones. *J Physiol* **509 ( Pt 1)**, 183-194 (1998). https://doi.org:10.1111/j.1469-7793.1998.183bo.x
